# Iron Chelation Improves Ineffective Erythropoiesis and Iron Overload in Myelodysplastic Syndrome Mice

**DOI:** 10.1101/2022.10.05.510967

**Authors:** Wenbin An, Maria Feola, Srinivas Aluri, Marc Ruiz-Martinez, Ashwin Shridharan, Maayan Levy, Eitan Fibach, Xiaofan Zhu, Amit Verma, Yelena Z. Ginzburg

**Affiliations:** Division of Hematology and Medical Oncology, Tisch Cancer Institute, Icahn School of Medicine at Mount Sinai, New York, NY; State Key Laboratory of Experimental Hematology, National Clinical Research Center for Blood Diseases, Division of Pediatric Blood Diseases Center, Institute of Hematology & Blood Diseases Hospital, Chinese Academy of Medical Sciences & Peking Union Medical College, Tianjin, 300020, China; Division of Hematology and Medical Oncology, Albert Einstein College of Medicine, Bronx, NY; Department of Hematology, Hadassah Medical Center, Hebrew University, Jerusalem, Israel

## Abstract

Myelodysplastic syndrome (MDS) is a heterogeneous group of bone marrow stem cell disorders characterized by ineffective hematopoiesis and cytopenias, most commonly anemia. Red cell transfusion therapy for anemia in MDS results in iron overload, correlating with reduced overall survival. Whether treatment of iron overload benefits MDS patients remains controversial. We evaluate underlying iron-related pathophysiology and the effect of iron chelation using deferiprone on erythropoiesis in *NUP98-HOXD13* transgenic mice, a highly penetrant well-established MDS mouse model. Our results characterize an iron overload phenotype with aberrant erythropoiesis in these mice which was reversed by deferiprone-treatment. Serum erythropoietin level decreased while erythroblast erythropoietin receptor expression increased in deferiprone-treated MDS mice. We demonstrate, for the first time, normalized expression of the iron chaperones *Pcbp1* and *Nco4* and increased ferritin stores in late stage erythroblasts from deferiprone-treated MDS mice, evidence of aberrant iron trafficking in MDS erythroblasts. Importantly, erythroblast ferritin is increased in response to deferiprone, correlating with decreased erythroblast ROS. Finally, we confirmed increased expression of genes involved in iron uptake, sensing, and trafficking in stem and progenitor cells from MDS patients. Taken together, our findings provide evidence that erythroblast-specific iron metabolism is a novel potential therapeutic target to reverse ineffective erythropoiesis in MDS.

**BRIEF SUMMARY:** Ineffective erythropoiesis in MDS mice correlates with aberrant iron trafficking within bone marrow erythroblasts, consistent with findings in MDS patient progenitors, reversed after iron chelation.

## INTRODUCTION

Myelodysplastic syndrome (MDS) is a heterogeneous group of bone marrow stem cell disorders characterized by ineffective hematopoiesis leading to blood cytopenias and increased incidence of transformation to acute myeloid leukemia (AML) [1,2]. In the US, MDS affects approximately 100,000 people with a median age of 65 years at diagnosis and an incidence of 40 / 100,000 per year thereafter [3,4]. Most MDS patients suffer from the accumulating consequences of marrow failure compounded by other age-related diseases. Several categories of MDS patients are low risk subtypes according to the revised International Prognostic Scoring System and have a longer median survival with the lowest rate of progression to AML [5,6]. Low risk MDS patients account for approximately two-thirds of all MDS patients with 30-50% requiring regular red blood cell (RBC) transfusions [7,8]. The main goals of therapy in low risk MDS patients are to alleviate cytopenias and their associated symptoms and thereby improve quality of life [9]. RBC transfusions remain the mainstay of therapy in low risk MDS [10] and are the main source of progressive iron overload and consequent end-organ damage in transfusion-dependent patients. Because of the low rates of progression to AML in low risk MDS patients, these patients have a substantial life expectancy and theoretically warrant screening for transfusional iron overload, known to increase morbidity and mortality in chronic RBC transfusion-dependent anemias [4,11,12]. Furthermore, although it is generally accepted that transfusional iron overload develops in low risk transfusion-dependent MDS patients and methods to diagnose and treat iron overload are available, the risk benefit ratio of treating iron overload in MDS patients remains controversial.

A correlation between iron overload and reduced survival has been demonstrated mostly by retrospective studies [13,14]. RBC transfusion-dependence correlates strongly with decreased survival in MDS patients [15,16], and MDS patients with elevated serum ferritin have significantly fewer Burst Forming Units (BFU-Es) but normal Granulocyte Macrophage Colony Forming Units (CFU-GM) [17]. Iron overload has been shown to inhibit BFU-E colony formation and erythroblast differentiation in both murine and human hematopoietic progenitors *in vitro* [18]. Finally, cells exposed to excess iron exhibit dysplastic changes with increased intracellular reactive oxygen species (ROS) and decreased expression of anti-apoptotic genes [18], and the addition of iron to MDS patients’ peripheral blood mononuclear cells resulted in increased ROS and DNA damage, triggering apoptosis [19,20]. These data suggest that iron overload may lead to a deleterious effect on hematopoiesis, worsening disease in MDS [21].

Along these lines, more recent studies demonstrate the potential benefit of iron chelation therapy on the overall survival in low risk MDS patients. Iron chelation is associated with improved hemoglobin (Hb) and reduced RBC transfusion requirements in some patients [10,22]. Most recently, the TELESTO trial demonstrated prolonged event-free survival in iron overloaded lower risk MDS patients treated with iron chelation (i.e. deferasirox) [23]. However, no clear improvement in Hb or reduction in RBC transfusions and no effect on overall survival was observed in deferasirox-treated MDS patients; this may be a consequence of the specific iron chelator selected. Mechanistically, iron chelation with deferiprone (DFP) results in increased hepcidin expression [24,25], changing the distribution of iron by moving it out of parenchymal cells and loading it onto circulating transferrin to enhance iron-mediated signaling to hepcidin expression. The relative affinities of iron chelators in comparison with transferrin determine whether the chelator can donate iron to transferrin [26]. As a consequence, only DFP, with a relatively lower iron binding capacity, can increase transferrin saturation and stimulate signaling to increase hepcidin expression in the liver. The physiologic effect of increased hepcidin in turn prevents further iron absorption and recycling, trapping iron within macrophages [27]. A direct beneficial effect of DFP on erythropoiesis in MDS has yet to be demonstrated.

Here, we evaluate the effect of iron chelation on erythropoiesis in NUP98-HOXD13 transgenic (NHD13) mice as a well-established *in vivo* model of MDS. We use NHD13 mice in light of the successful pre-clinical use of this model to investigate the effects of an activin receptor II ligand trap on reversing ineffective erythropoiesis in MDS mice [28]. We also use the iron chelator DFP in light of its ability to cross the cell membrane and mobilize intracellular iron and its ability to donate iron to transferrin [26]. Our results demonstrate that NHD13 mice exhibits anemia, increased serum erythropoietin (EPO), expanded erythropoiesis in the bone marrow and spleen, and parenchymal iron overload, consistent with a low risk MDS phenotype in patients. In addition, we demonstrate decreased EPO-responsiveness, decreased erythroblast differentiation, and impaired enucleation in bone marrow erythroblasts from MDS mice. Furthermore, iron chelation with DFP, in addition to decreasing parenchymal iron deposition, restores hepdicin:iron responsiveness, partially reverses anemia, and normalizes serum EPO concentration in MDS mice. DFP-treated MDS mice also exhibit normalized erythroblast differentiation, expression of *Gata1* and *Epor* (EPO receptor) as well as that of iron chaperones *Pcbp1* and *Ncoa4*, and increased erythroblast ferritin concentration in bone marrow erythroblasts. Finally, we demonstrate aberrant expression of genes involved in iron uptake, sensing, and trafficking in MDS patient bone marrow stem and progenitor cells. Taken together, our data for the first time provides *in vivo* evidence that ineffective erythropoiesis in MDS mice is responsive to iron chelation with DFP, normalizing erythroblast iron trafficking and restoring EPO responsiveness to reverse anemia in MDS.

## RESULTS

### MDS Mice Exhibit Elevated MCV Anemia with Expanded Erythropoiesis and Iron Overload, Appropriate as a Model of Low Risk MDS

To establish the expected MDS phenotype, mice were sacrificed at 6 months of age and demonstrate significantly reduced red blood cell (RBC) counts and Hb concentration, increased MCV, and no difference in reticulocyte count relative to WT controls (Table I). MDS mice also exhibit decreased white blood cell (WBC) count and no difference in platelet count (Table I). Consistently, our experiments also demonstrate that MDS mice exhibit significantly increased serum EPO concentration and increased bone marrow cellularity (Table I). Furthermore, we demonstrate borderline increased apoptosis in bone marrow erythroblast in MDS mice, reaching significance in OrthoE (Supplementary Figure 1). Finally, liver iron concentration is significantly increased in MDS mice (Table I). In conjunction with previously published work [28-30], this combination of characteristics provides substantial evidence that NHD13 mice are an appropriate mouse model of MDS [31].

**Table I:** Characteristics of NHD13 mice are consistent with elevated MCV anemia with expanded hematopoiesis and iron overload which is appropriate as a mouse model of MDS.

|  | RBC<br>count | Hb | MCV | Retic<br>count | WBC<br>count | Platelet<br>count | Serum<br>EPO | Bone<br>marrow<br>cells | Liver iron<br>concentration |
| --- | --- | --- | --- | --- | --- | --- | --- | --- | --- |
| (units) | (10<sup>6</sup>/μL) | (g/dL) | (fL) | (10<sup>6</sup>/μL) | (10<sup>6</sup>/μL) | (10<sup>3</sup>/μL) | (μg/μL) | (10<sup>7</sup> cells) | (mg/g dry<br>weight) |
| **WT** | 10±0.1 | 14.7±0.1 | 50±0.5 | 495±67 | 5.5±0.6 | 759±67 | 308±48 | 12.4±0.7 | 0.23±0.07 |
| **MDS** | 6.73±0.29 | 11.2±0.4 | 60±1.3 | 440±30 | 3.0±0.4 | 672±80 | 4832<br>±1154 | 15±0.8 | 0.51±0.04 |
|  | *** | *** | *** | NS | ** | NS | *** | * | * |

WT = wild type; MDS = myelodysplastic syndrome (NHD13) mice; RBC = red blood cell; Hb = hemoglobin; MCV = mean corpuscular hemoglobin; Retic = reticulocyte; WBC = white blood cell; EPO = erythropoietin; NS = not significant; \* $P<0.05$ ; \*\* $P<0.01$ ; \*\*\* $P<0.0001$ .

### Iron Chelation with DFP Reverses Iron Overload, Normalized Erythroferrone Expression in Bone Marrow Erythroblasts, and Improves Hepcidin Iron Responsiveness in MDS Mice

Next, we evaluate the effects of iron chelation with DFP in MDS mice. First, we demonstrate that DFP can be detected in the serum of DFP-treated mice (Supplementary Figure 2). Second, our results demonstrate that DFP-treated MDS mice exhibit increased serum iron concentration and transferrin saturation (Figure 1A and 1B); both male and female mice demonstrate equivalent responses to DFP (data not shown). These findings are consistent with reversal of increased parenchymal iron loading in MDS mice after DFP treatment, exhibiting decreased liver, spleen, and bone marrow non-heme iron concentration (Figure 1C-1E) and validate the expected re-distribution of iron from parenchymal deposition to the circulating compartment, to ultimately enable excretion. Furthermore, ferritin concentration is not statistically significantly different in MDS bone marrow erythroblasts but increased in DFP-treated MDS bone marrow erythroblasts (Supplementary Figure 3A and 3B). Finally, *Hamp* expression, the gene encoding for hepcidin, in the liver is unchanged and *Hamp*-iron responsiveness is decreased in MDS relative to WT mice (Figure 1F and 1G). While *Hamp* expression is not increased in the liver of DFP-treated MDS mice (Figure 1F), it is significantly increased relative to non-heme iron concentration in the liver (Figure 1G) providing evidence of enhanced hepcidin responsiveness to iron in DFP-treated MDS mice.

**Figure 1.**
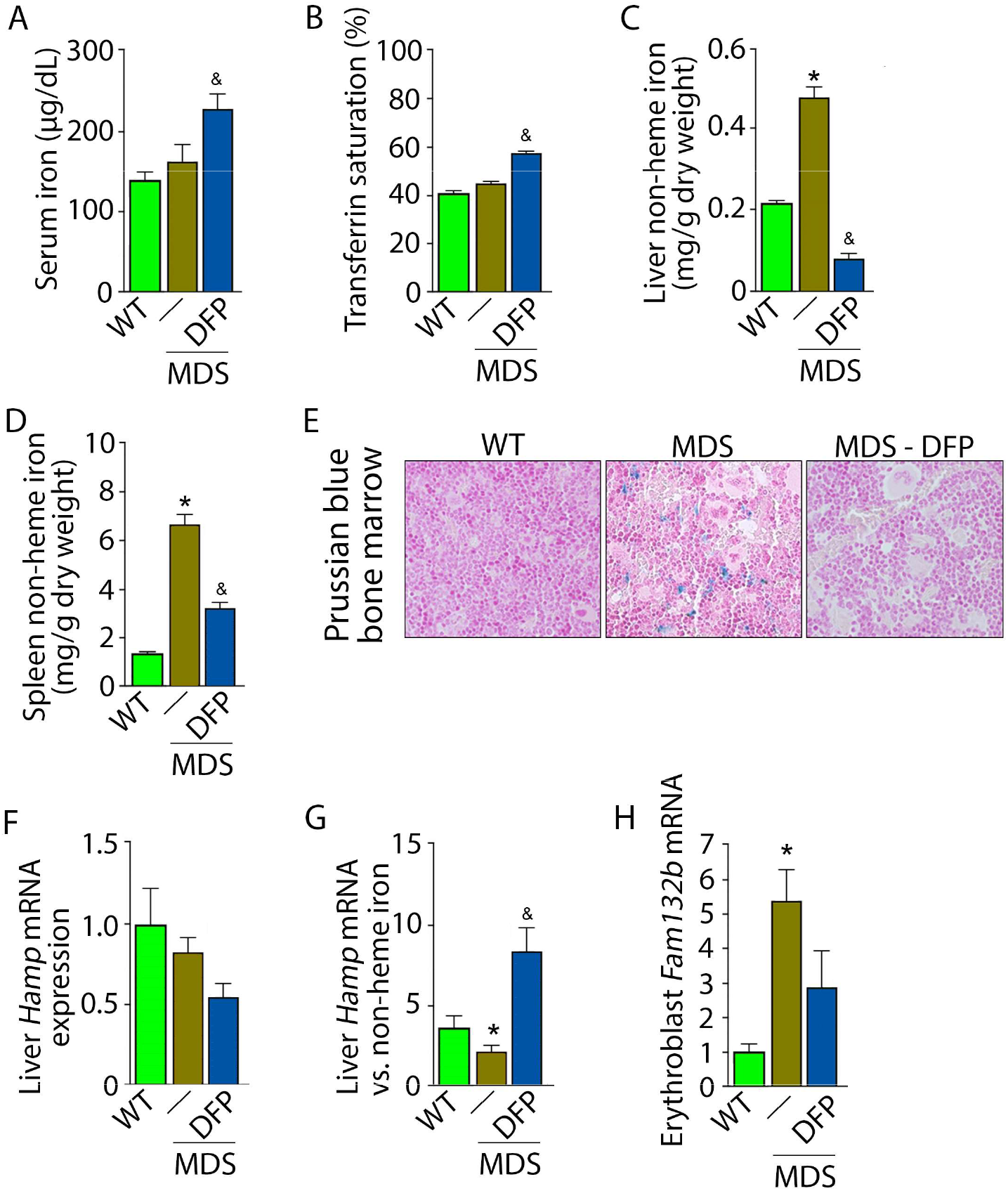
DFP reverses parenchymal iron overload and restores hepcidin iron responsiveness in MDS mice. DFP results in increased serum iron **(a)** and transferrin saturation **(b)** while reducing parenchymal iron in the liver, spleen, and bone marrow **(c-e)**, While liver *Hamp* mRNA expression is unchanged in WT, MDS, and DFP-treated MDS mice **(f)**, *Hamp* responsiveness to iron is normalized in DFP-treated MDS mice **(g)** (n = 7-10 mice/group). **(h)** DFP results in more normal *Fam132b* mRNA expression (n = 10-12 mice/group) in sorted bone marrow erythroblasts from MDS mice analyzed after 1 month of treatment. \**P*< 0.05 vs. WT; ^&^*p* < 0.05 vs. MDS; Abbreviations: WT = wild type; MDS = myelodysplastic syndrome; DFP = deferiprone; *Hamp* = hepcidin; *Fam132b* = erythroferrone.

Hepcidin expression represents the effect of multiple pathways. Specifically, hepcidin is upregulated in response to liver iron stores and circulating iron; increased in the setting of inflammation; and downregulated in conditions of expanded or ineffective erythropoiesis as a consequence of elevated *Fam132b* expression, the gene name for erythroferrone (ERFE), in erythroblasts [32]. Our results demonstrate that DFP-treated MDS mice exhibit decreased liver iron concentration (Figure 1C) while increasing transferrin saturation (Figure 1B) and no evidence of inflammation-mediated STAT3 signaling to hepcidin in DFP-treated MDS mouse liver (Supplementary Figure 4A and 4B). Thus, the effect of iron redistribution on hepcidin expression in DFP-treated MDS mice is predictably small, leading us to also evaluate the contribution of changes in erythropoiesis on increased hepcidin responsiveness in DFP-treated MDS mice.

Erythroblast *Fam132b* expression is increased in MDS mouse bone marrow (Figure 1H), the response expected in the setting of increased serum EPO concentration [33] in MDS relative to WT mice. Similar to other diseases of ineffective erythropoiesis, increased expression of bone marrow *Fam132b* is expected to suppress hepcidin and decrease hepcidin iron responsiveness, resulting in iron overload [33,34]. As a consequence, our current findings demonstrate that increased bone marrow erythroblast *Fam132b* expression results in inappropriately low liver *Hamp* expression relative to parenchymal iron loading (Figure 1C-1G), demonstrating decreased *Hamp* responsiveness to iron in MDS mice. These findings further support use of these mice as an appropriate MDS model in which decreased hepcidin iron responsiveness is a consequence of expanded erythropoiesis, leading to systemic iron overload observed in this disease [31]. Furthermore, reversal of ineffective erythropoiesis in DFP-treated MDS mice, with normalization of *Fam132b* expression (Figure 1H), leads to restored *Hamp* responsiveness to iron and reversal of parenchymal iron loading in the liver, spleen, and bone marrow (Figure 1C-1G). Finally, erythroblast *Fam132b* expression is not altered in DFP-treated WT mice (Supplementary Figure 5). Taken together, our results provide further evidence that mitigating the ERFE hepcidin pathway is central to the pathophysiology of ineffective erythropoiesis and its reversal.

### Iron Chelation with DFP Improves Ineffective Erythropoiesis in MDS Mice

Five month old MDS mice were treated for 4 weeks with DFP and sacrificed for analyses at 6–months of age. DFP-treated MDS mice exhibit increased Hb and RBC count relative to untreated MDS mice and no change in MCV or reticulocytosis (Figure 2A-2D); both male and female mice demonstrate equivalent responses to DFP (data not shown). In addition, the WBC count remained low while platelet and neutrophil counts are unchanged in DFP-treated relative to untreated MDS mice (Figure 2E-2G). Furthermore, while the spleen size increased in MDS relative to WT mice and is not significantly decreased after DFP (Figure 3A), splenic architecture is improved (Figure 3B)—with relatively decreased red pulp and more organized splenic nodules—and serum EPO concentration is normalized (Figure 3C) in DFP-treated relative to untreated MDS mice. Consistently, the total number of erythroblasts and the erythroid fraction in the bone marrow are increased in MDS relative to WT mice and normalized in DFP-treated relative to untreated MDS mice, evident decrease especially in BasoE and PolyE fractions (Figure 3D-3F). DFP also leads to a normalized bone marrow erythroblast differentiation in MDS mice (Figure 3G) with a proportionally increased PolyE and decreased OrthoE fractions in bone marrow from MDS mice, normalized after DFP treatment, consistent with a block of erythroblast differentiation at PolyE in MDS patients [35]. Finally, erythroblast apoptosis is unchanged (Figure 3H) despite decreased erythroblast ROS (Figure 3I) in DFP-treated relative to untreated MDS mice. These findings are globally consistent with improvement in ineffective erythropoiesis in response to DFP treatment without effects on erythroblast apoptosis in MDS.

**Figure 2.**
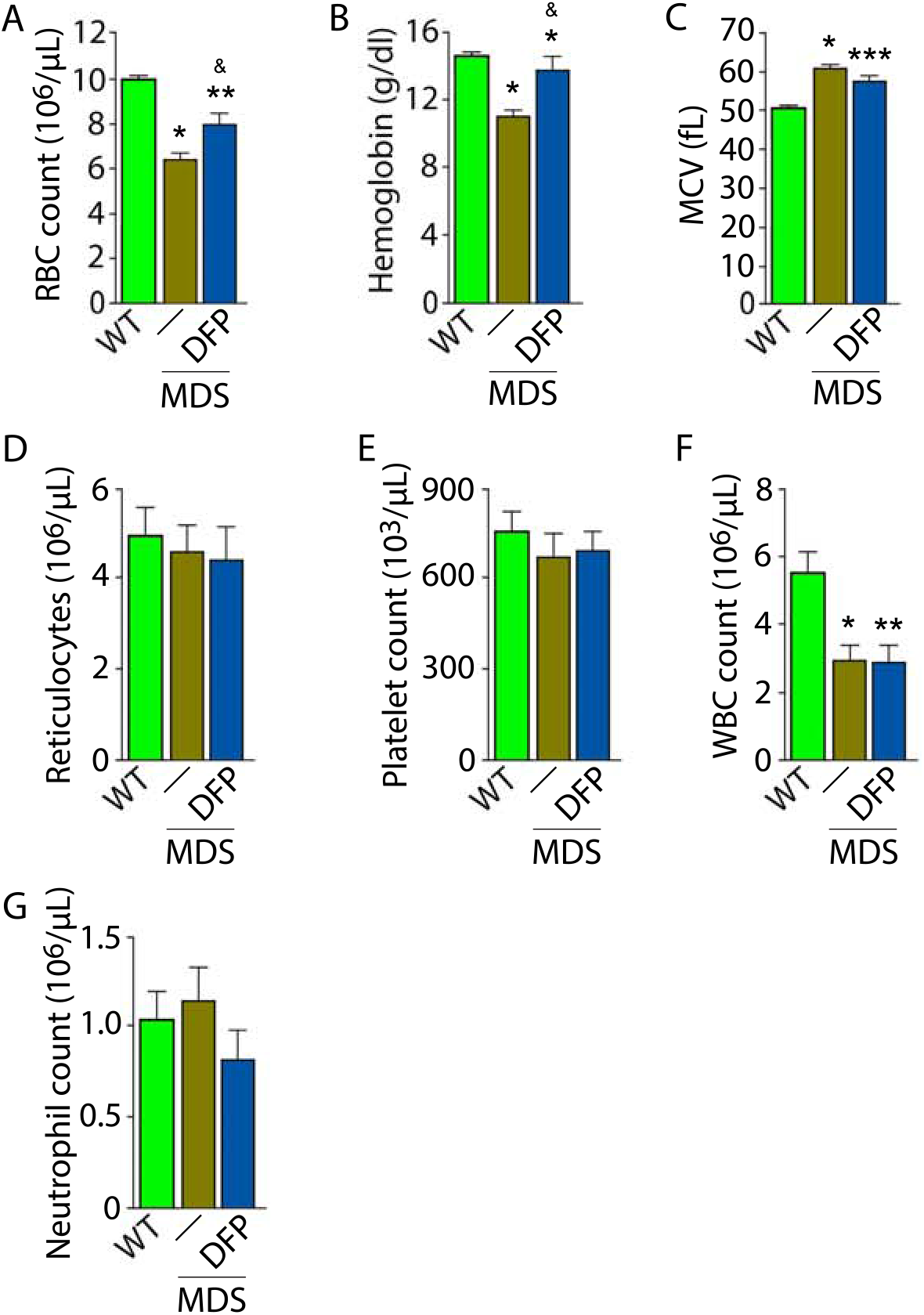
Elevated MCV anemia in MDS mice is partially reversed by DFP. Circulating RBC count **(a)**, hemoglobin **(b)**, MCV **(c)**, reticulocyte count **(d)**, platelet count **(e)**, WBC count **(f)**, and neutrophil count **(g)** in WT, MDS, and DFP-treated MDS mice (n = 10-14 mice/group) analyzed after 1 month of treatment. \**P*<0.05 vs. WT; ** *P*<0.01 vs. WT; *** *P*<0.001 VS. WT; ^&^*P*<0.05 VS. MDS; Abbreviations: WT= wild type; MDS= myelodysplastic syndrome; DFP= deferiprone; RBC = red blood cell; MCV = mean corpuscular volume; WBC = white blood cell.

**Figure 3.**
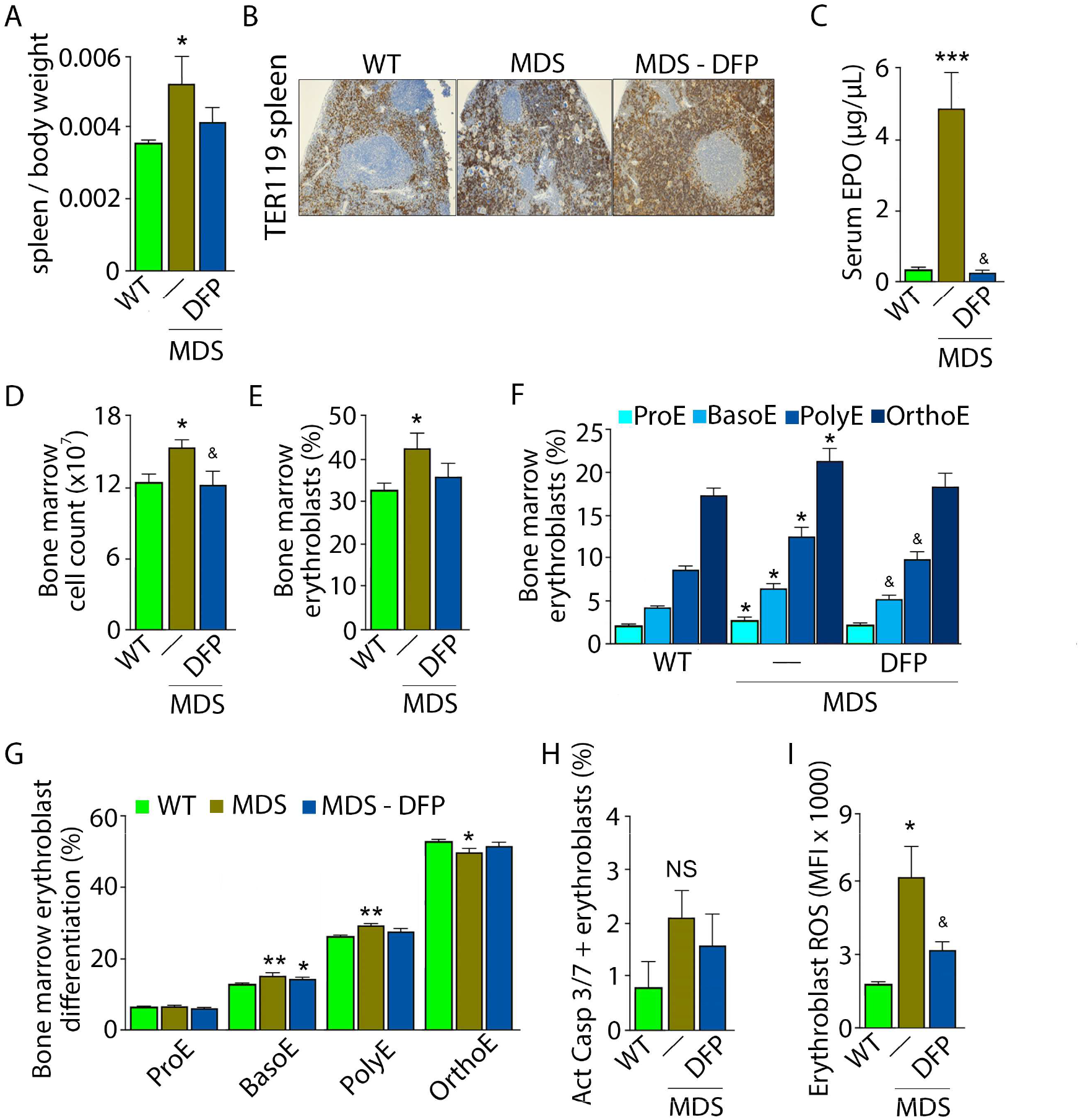
Expanded and ineffective erythropoiesis in MDS mice is partially reversed by DFP. Spleenweight (n = 11-12 mice/group) **(a)**, splenic architecture (n = 5 mice/group) **(b)**, serum EPO concentration (n = 5-12 mice/group) **(c)**, and bone marrow erythroblast count (n = 13-15 mice/group) **(d)** are more normal in DFP-treated MDS mice analyzed after 1 month of treatment. The total fraction of erythroblasts in the bone marrow is increased in MDS relative to WT mice, with no difference between WT and DFP-treated MDS mice (n = 13-15 mice/group) **(e)**. The fraction of all stages of terminal erythropoiesis is increased in MDS relative to WT mice and decreased in DFP-treated relative to untreated MDS mice in BasoE and PolyE stages (n = 13-15 mice/group) **(f)**. Erythroblast differentiation in the bone marrow, decreased in MDS relative to WT, is normalized in DFP-treated relative to untreated MDS mice (n = 13-15 mice/group) **(g)**. In addition, erythroblast apoptosis, as measured by activated caspase 3/7, is unchanged in DFP treated MDS mice (n = 7-11 mice/group) **(h)**.Finally, erythroblast ROS is decrease in DFP-treated relative to untreated MDS mice (n = 11-12 mice/group) **(i)** analyzed after 1 month of treatment. \**P*<0.05 vs. WT; ** *P*<0.01 vs. WT; *** *P*<0.001 vs. WT; ^&^*P*<0.05 vs. MDS; Abbreviations: WT= wild type; MDS = myelodysplastic syndrome; DFP = deferiprone; EPO = erythropoietin; Act casp *3/7* = activated caspase 3 and 7; ROS = reactive oxygen species; ProE = pro-erythroblasts; BasoE = basophilic erythroblasts; PolyE = polychromatophilic erythroblasts; OrthoE = orthochrom atophiIic erythroblasts; Ns = not significant.

Importantly, we evaluate the effects of DFP in WT mice. Specifically, our results demonstrate no change in RBC count, Hb, MCV, reticulocyte count, or serum EPO in DFP-treated relative to untreated WT mice (Supplementary Figure 6A-6E). In addition, the bone marrow erythroblast fraction in DFP-treated WT mice is decreased, especially in the late stages of terminal erythropoiesis (i.e. PolyE and OrthoE), similar to DFP-treated MDS mice. However, unlike DFP-treated MDS mice, erythroid differentiation is decreased in DFP-treated relative to untreated WT mice (Supplementary Figure 6F-6H). Furthermore, similar to DFP-treated MDS mice, DFP in WT mice results in unchanged erythroblast apoptosis (Supplementary Figure 6I) albeit without affecting ROS (Supplementary Figure 6J). Taken together, these findings support our conclusions that reversal of ineffective erythropoiesis in MDS mice occurs independently of changes in erythroblast apoptosis, that DFP has a direct effect on erythropoiesis, and that the differences in effect on MDS and WT mice are dependent on the effectiveness of erythropoiesis in the underlying state. Finally, we confirmed that the effect of DFP in WT mice occurs despite a relatively higher serum DFP concentration and metabolized DFP-G concentration in WT relative to MDS mice (Supplementary Figure 7).

### Normalized Expression of EPO Downstream Genes in Bone Marrow Erythroblast from DFP-treated MDS Mice

To evaluate erythropoiesis more closely, we also measure *Gata1* and *Bcl-Xl* expression to assess effect of DFP on other EPO-STAT5 target genes in MDS erythroblasts. *Gata1* and *Bcl-Xl* expression is known to be downstream of EPO. While GATA1 is as an important transcriptional regulator in normal erythropoiesis [36], BCL-xl is implicated in the anti-apoptosis effect of EPO on erythroblasts [37]. Aberrant GATA1 expression in MDS have been described with evidence of increased GATA1 expression in bone marrow CD34+ stem and progenitor cells as well as CD71+ erythroblasts from MDS patients [38]. Furthermore, normal upregulation of GATA1 and BCL-xl during human erythroid differentiation is lost in MDS [39]. Neither *Gata1* or *Bcl-Xl* expression have previously been evaluated in MDS mice; their expression is expected to increase in conditions of elevated EPO concentration.

Our results demonstrate that *Gata1* mRNA expression is increased in sorted bone marrow ProE, borderline decreased in BasoE, and decreased in PolyE and OrthoE erythroblasts from MDS relative to WT mice (Figure 4A-4D), consistent with expectations that GATA1 expression is elevated in MDS patient bone marrow stem and progenitor and early erythroblasts [38] with loss of upregulation during erythroblast differentiation [39]. DFP treatment restores *Gata1* mRNA expression relative to untreated MDS or WT mice (Figure 4A-4D). Furthermore, *Bcl-Xl* mRNA expression is decreased in bone marrow erythroblasts from MDS relative to WT mice and return to normal expression levels in DFP-treated MDS mice (Figure 4F and 4G) despite increased serum EPO (Figure 3C) and borderline increased erythroblast apoptosis (Figure 3H) in MDS erythroblasts, normalized in DFP-treated relative to untreated MDS mice. Bone marrow erythroblast *Bcl-Xl* expression is also increased in DFP-treated WT mice (Supplementary Figure 8). These results raise an important question, namely whether physiological or pathophysiological nuances in EPO-STAT5 signaling can conceptually separate EPO responsiveness from EPO mediated anti-apoptotic effects in erythroblasts.

**Figure 4.**
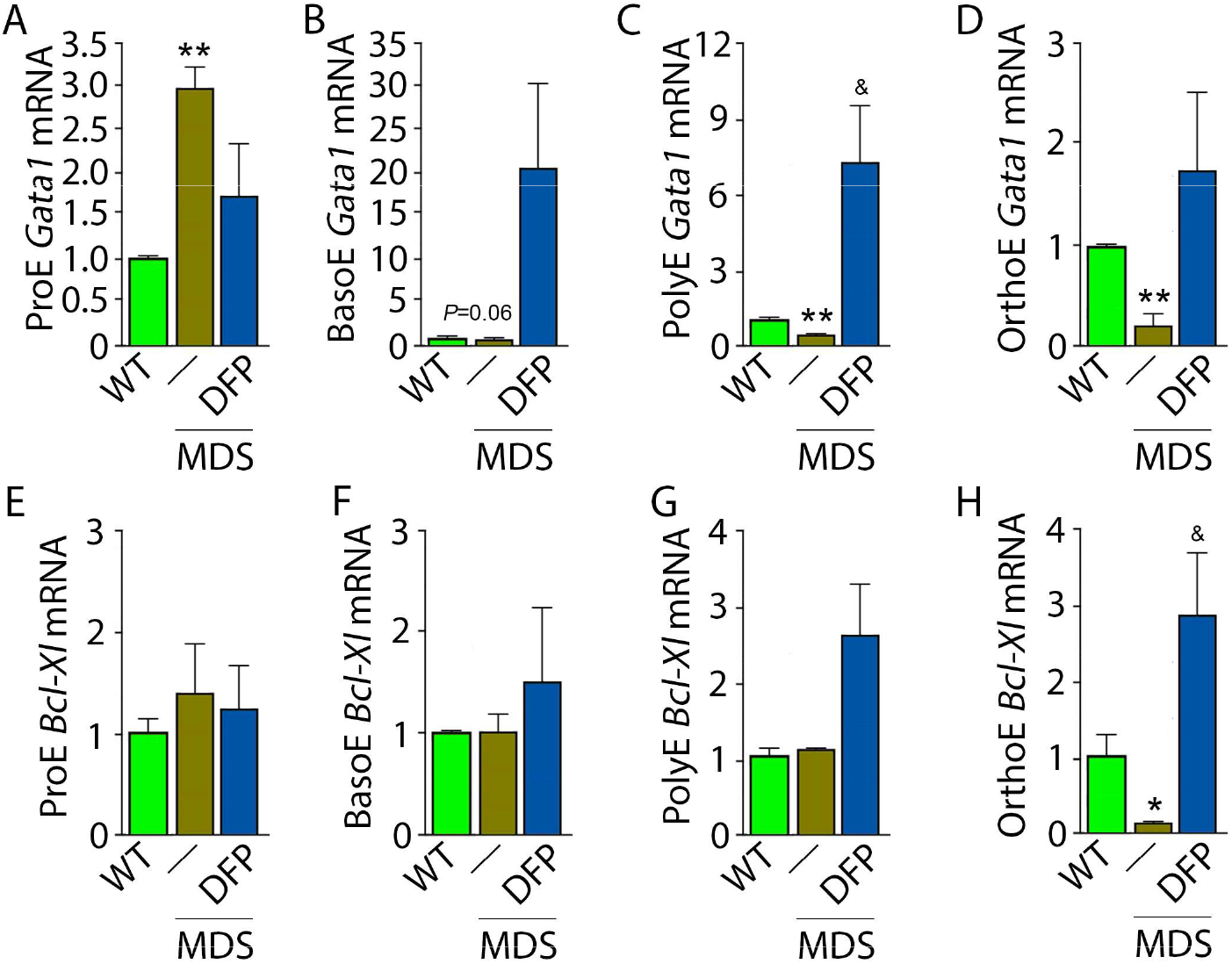
DFP leads to normalized gene expression downstream of EPO in MDS erythroblasts. *Gata1* mRNA expression is increased in sorted bone marrow ProE **(a)**, borderline decreased in BasoE **(b)**, and significantly decreased in PolyE **(c)** and OrtlloE **(d)** erythroblasts from MDS relative to WT mice: DFP treatment restores *Gata1* mRNA expression relative to untreated MDS or WT mice (n = 15-21 mice/group). DFP treatment in MDS mice does not affect *Bcl-Xl*mRNA expression in sorted bone marrow ProE **(e)**, BasoE **(f)**, and PolyE **(g)**, and increases it in OrthoE **(h)** erythroblasts relative to untreated MDS or WT mice (n = 10-12 mice/group). \**P*<0.05 VS. WT; \*\**P*<0.01 VS. WT; ^&^*P*<0.05 VS. MDS; Abbreviations: WT = Wild type; MDS = myelodysplastic syndrome; DFP = deferiprone: Gata1 = erythroid transcription factor. Bcl-XI = B-cell lymphoma-extra large: ProE = pro-erythroblasts; BasoE = basophilic erythroblasts; PolyE = polychromatophilic erythroblasts: OrthoE = orthochromatophilic erythroblasts.

We then evaluate signaling pathways downstream of EPO. Both STAT5 and AKT signaling are essential for erythropoiesis. Prior work demonstrates that the expected STAT5 signaling response to EPO is hampered by iron restriction [40]. Others demonstrate that AKT signaling is implicated in EPO-mediated erythroblast survival [41], essential in conditions with elevated EPO when *Epor* expression is suppressed [42-44]. We demonstrate enhanced STAT5 and AKT phosphorylation in MDS relative to WT bone marrow erythroblasts without changes in erythroblasts from DFP-treated relative to untreated MDS mice (Supplementary Figure 9). These findings suggest that the expected changes in signaling downstream of EPO are unaffected by DFP administration, implicating possible changes in erythroblast *Epor* expression in DFP-treated MDS mice.

### DFP Increases *Epor* Expression in Later Stage MDS Erythroblasts

Next, we explore erythroblast *Epor* expression in DFP-treated and untreated MDS mice. We hypothesize that *Epor* plays an important role in EPO responsiveness that is independent of EPO concentration. This hypothesis is based on findings in EpoR-H mice, a knock-in mutation leading to normal EPO-EpoR binding and signaling but absent EpoR internalization and degradation [45-47]. These mice exhibit decreased serum EPO levels, elevated RBC counts, and a smaller proportion of mature erythroid precursors in the bone marrow relative to WT mice [40], suggesting that EpoR expression may influence erythroblast differentiation in a manner that is complementary to the anti-apoptotic effect of EPO.

Based on this premise, we anticipate that *Epor* expression is decreased in bone marrow erythroblast from MDS relative to WT mice, restored in DFP-treated relative to untreated MDS mice. We used sorted bone marrow to evaluate *Epor* expression in progressive stages of terminal erythropoiesis. First, *Epor* expression is borderline increased in ProE and significantly increased in BasoE, earlier stage erythroblasts (Figure 5A and 5B) and decreased in PolyE and OrthoE, later stage erythroblasts (Figure 5C and 5D), in MDS relative to WT mice. Second, *Epor* expression is significantly increased in all stage erythroblasts in DFP-treated relative to untreated MDS or WT mice (Figure 5A-5D). Finally, DFP does not result in increased *Epor* expression in bone marrow erythroblasts from WT mice (Supplementary Figure 10). Taken together, increased later stage erythroblast *Epor* expression is a potential mechanism by which DFP leads to enhanced EPO responsiveness and enhanced erythroid differentiation despite decreased serum EPO concentration, reversing ineffective erythropoiesis exclusively in MDS mice.

**Figure 5.**
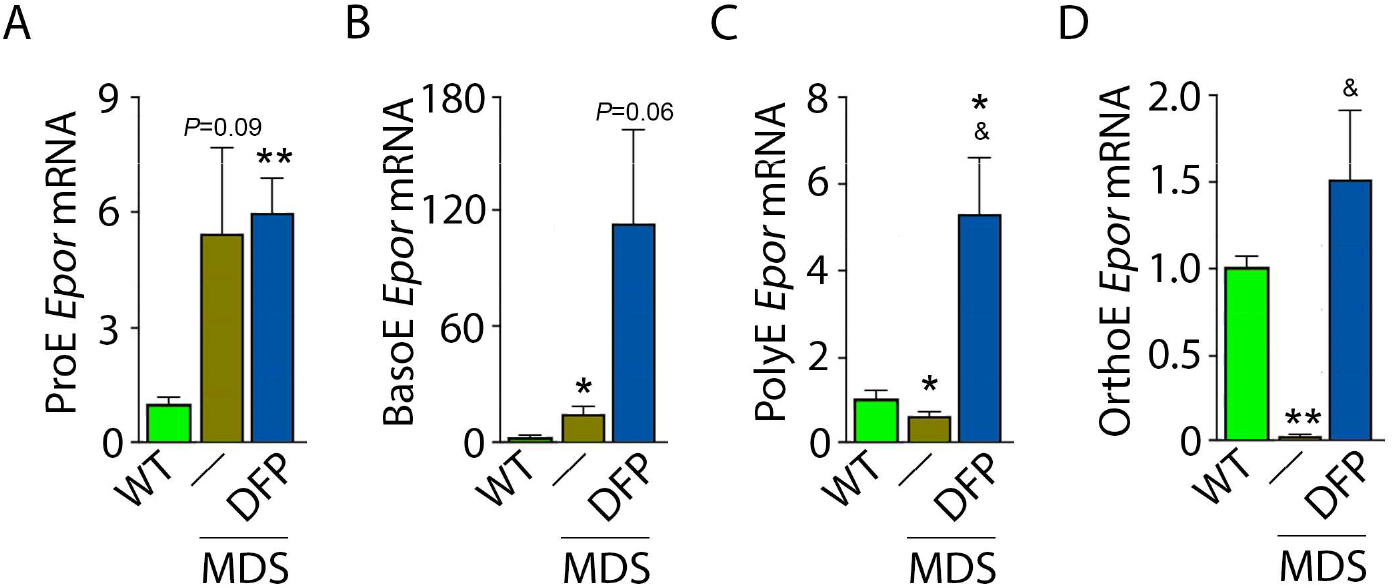
DFP increases cporexpression in MDS erythroblasts. DFP treatment in MDS mice increases *Epor* mRNA expression in bone marrow ProE **(a)**, BasoE **(b)**, PolyE **(c)**, and OrthoE erythroblasts relative to untreated MDS or WT mice. **(e)** When *Epor* mRNA expression is relativized to that in ProE, DFP leads to relatively higher *Epor* expression in PolyE and OrthoE stage erythroblasts from MDS mice (n = 15-18 mice / group). \**P*<0.05 vs. WT; * \**P*<0.01 vs. WT; ^&^*P*<0.05 ^VS.^ MDS; Abbreviations: WT = wild type; MDS = myelodysplastic syndrome; DFP = deferiprone; Epor = erythropoietin receptor; ProE = pro-erythroblasts; BasoE = basophilic erythroblasts; P olyE = polychromatophilic erythroblasts; OrthoE = orthochromatophilic erythroblasts.

### Defective Enucleation in MDS Erythroblasts is Normalized by DFP

Because increased EPO is implicated in defective enucleation [44], we also evaluate erythroblast enucleation in WT, MDS, and DFP-treated MDS mice. Our results demonstrate decreased enucleation in bone marrow erythroblasts from MDS relative to WT mice and return to normal expression levels in DFP-treated MDS mice (Supplementary Figure 11A and 11B). These findings are consistent with prior work which provide mechanistic evidence of an enucleation defect in MDS [48]. Our findings are also consistent with our previously published evidence demonstrating that manipulating iron metabolism in erythropoiesis leads to reversal of ineffective erythropoiesis is associated with normalization of the erythroblast enucleation defect in β-thalassemic mice, another model of ineffective erythropoiesis [49].

### TFR1 but not TFR2 Expression in MDS Erythroblasts is Normalized by DFP

Decreased iron in the bone marrow of DFP-treated MDS mice prompted us to evaluate whether specific mechanisms involved in iron sensing and trafficking could explain the beneficial effects of DFP on ineffective erythropoiesis in MDS mice. First, we demonstrate increased cell surface TFR1 on bone marrow erythroblasts from MDS relative to WT mice, normalized in DFP-treated MDS mice (Figure 6A); this pattern of erythroblast surface TFR1 is replicated in all stages of terminal erythropoiesis (data not shown). Similarly, cell surface TFR1 on bone marrow erythroblasts from DFP-treated WT mice is decreased relative to WT mice (Supplementary Figure 12). Because *Tfr1* expression in bone marrow is mainly EPO-mediated [50], we anticipate that erythroblast *Tfr1* expression is elevated in MDS as a consequence of high EPO, consequently decreased in DFP-treated relative to untreated MDS mice. We used sorted bone marrow to evaluate *Tfr1* expression in progressive stages of terminal erythropoiesis. First, *Tfr1* expression is increased in ProE and BasoE stages in MDS mice and remains elevated in DFP-treated MDS mice (Figure 6B and 6C). Next, *Tfr1* expression is normal and decreased in PolyE and OrthoE stages, respectively, in MDS relative to WT mice and increased to normal levels in DFP-treated relative to untreated MDS mice (Figure 6D and 6E). These findings demonstrate that *Tfr1* expression correlates with *Epor* expression while serum EPO remains the primary determinant of erythroblast cell surface TFR1, further coordinating EPO-responsiveness and iron uptake during erythropoiesis.

**Figure 6.**
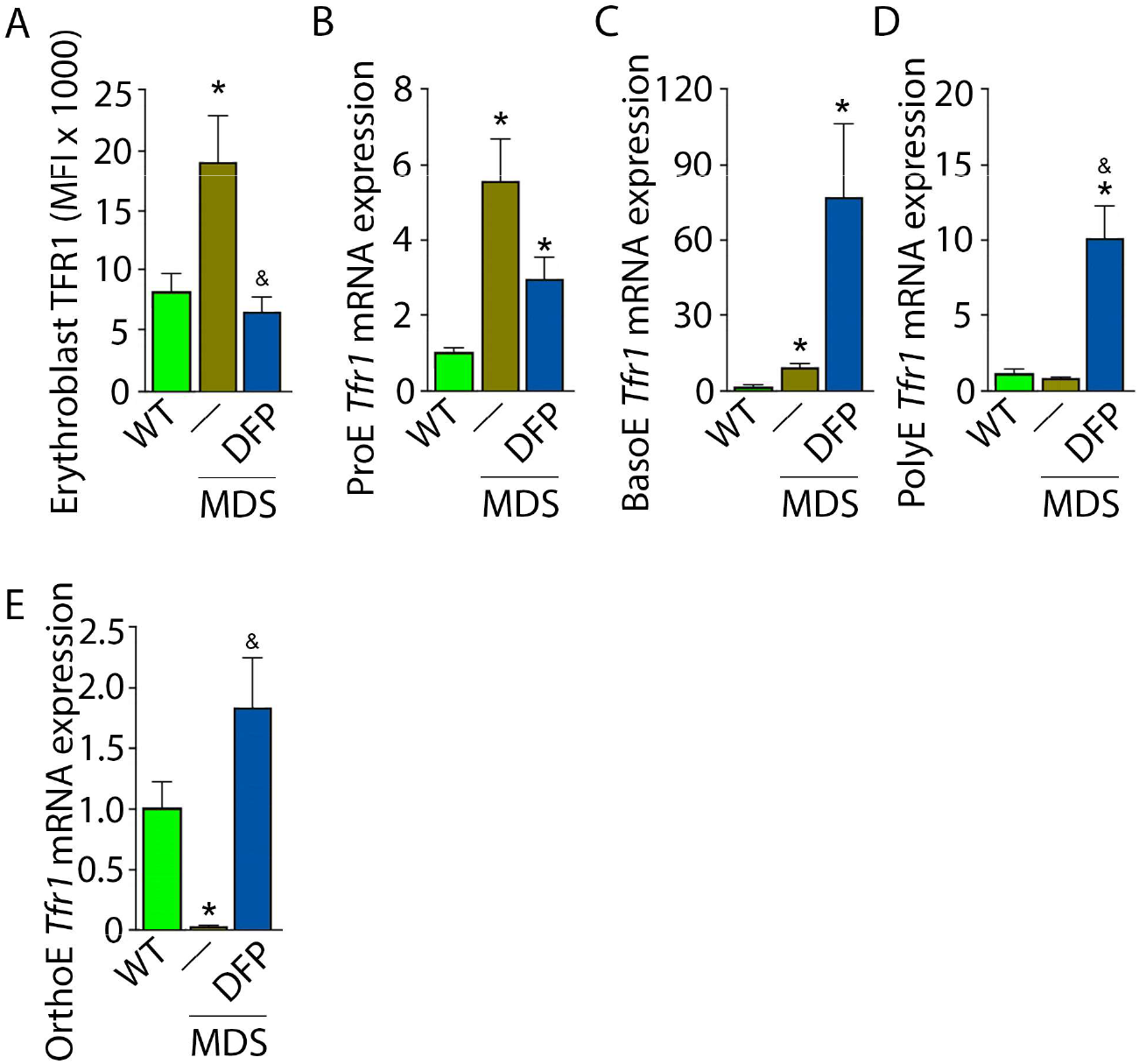
DFP nom alized TFR1 expression in MDS erythroblasts. **(a)** Bone marrow erythroblast membrane TFR1 is increased in MDS mice, normalized in response to DFP (n = 7-13 mice/group) analyzed after 1 month of treatment. *Tfr1* mRNA expression is increased in sorted bone marrow ProE **(b)** and BasoE **(c)** from MDS mice and remains elevated in DFP-treated MDS mice. *Tfr1* mRNA expression is normal in sorted bone marrow PolyE **(d)** and decreased in OrthoE and increased in OFP-treated relative to untreated MDS mice (n = 15-18 mice/group). \**P*<0.05 vs. WT; ^&^*P*<0.05 VS. MDS; Abbreviations: WT= wild type; MDS = myelodysplastic syndrome; DFP = deferiprone; TFR1 = transferrin receptor 1; ProE = pro-erythroblasts; BasoE = basophilic erythroblasts; PolyE = polychromatophilic erythroblasts; OrthoE = orthochromatophilic erythroblasts.

Next, we evaluate levels of TFR2 in light of its role in iron sensing and coordination of EPO-responsiveness with iron supply during erythropoiesis [40,51]. Furthermore, TFR2 is under investigation as a potential therapeutic target in β-thalassemia, another disease of ineffective erythropoiesis [52]. We hypothesize that TFR2, given the proposed interaction with EPOR [53], plays a central compensatory role in ineffective erythropoiesis. MDS mice exhibit higher erythroblast surface TFR2 expression specifically in ProE relative to WT, unchanged in DFP-treated MDS mice (Figure 7A). In addition, *Tfr2* expression is also borderline increased in bone marrow ProE erythroblasts, significantly higher in DFP-treated MDS relative to WT mice (Figure 7B); no significant differences are evident in BasoE, PolyE, and OrthoE bone marrow erythroblasts between WT, MDS, and DFP-treated MDS mice (Supplementary Figure 13A-13C). When compared with ProE, *Tfr2* expression in BasoE, PolyE, and OrthoE is significantly suppressed and remains suppressed in DFP-treated MDS mice (Figure 7C). Finally, no obvious differences in TFR2 protein concentration (Figure 7D and 7E) or *Scrib* mRNA expression (Supplementary Figure 14A-14D) are evident between bone marrow erythroblasts from WT, MDS, and DFP-treated MDS mice. These findings do not identify a DFP-specific modification of TFR2 mRNA, protein, or erythroblast surface localization or its action through changes in *Scrib* [40]. A mechanistic role for TFR2 in ineffective erythropoiesis in MDS remains to be fully elucidated.

**Figure 7.**
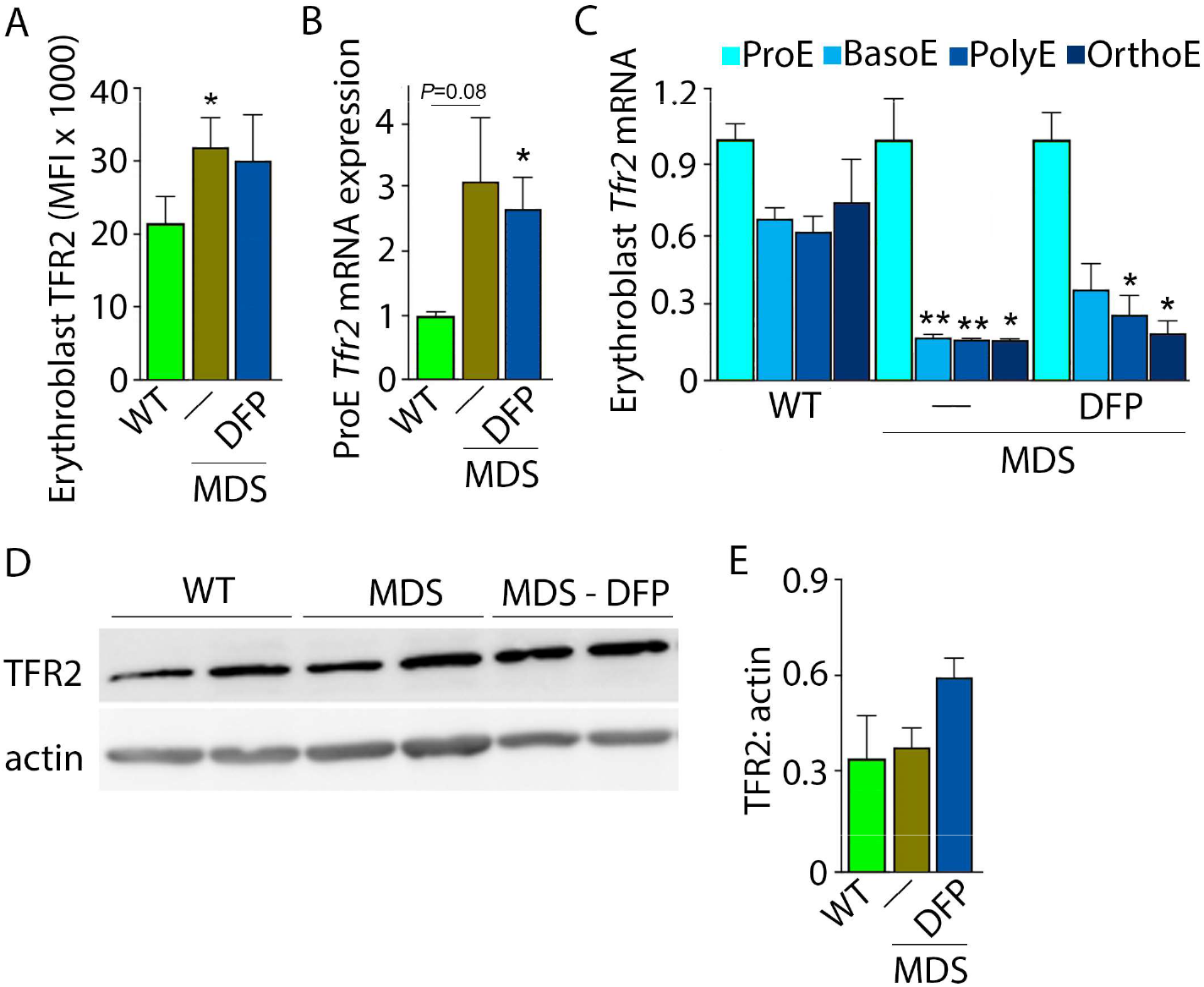
DFP does not alter TFR2 expression in MDS erythroblasts. **(a)** Bone marrow erythroblast membrane TFR2 is increased in MDS mice and remains elevated in DFP-treated MDS mice (n = 6-7 mice/group) analyzed after 1 month of treatment. **(b)** *Tfr2* mRNA expression is borderline increased in sorted bone marrow ProE from MDS relative to WT mice and remains significantly elevated in DFP-treated MDS relative to WT mice (n = 15-18 mice/group). **(c)** When compared with sorted bone marrow ProE, *Tfr2* expression in sorted bone marrow BasoE, PolyE, and OrthoE is significantly suppressed in MDS relative to WT mice and remains suppressed in DFP-treated MD s mice (n = 15-18 mice/group). **(d)** TFR2 protein concentration in bone marrow erythroblasts is not different between WT, MDS, and DFP-treated MDS mice, quantified in **(e)** (n = 3 mice/group). \**P*<0.05 vs. WT; \*\**P*<0.01 vs. WT; Abbreviations: WT = wild type; MDS = myelodysplastic syndrome; DFP = deferiprone; TFR2 = transfenin receptor 2; ProE = pro erythroblasts; BasoE **=** basophilic erythroblasts; PolyE **=** polychromatophilic erythroblasts; OrthoE **=** orthochromatophilic erythroblasts.

### Altered Iron Trafficking in MDS Erythroblasts is Partially Restored by DFP

We explored further the expression of other iron chaperones in MDS mouse bone marrow erythroblasts, hypothesizing that improved erythropoiesis in DFP-treated MDS mice is a consequence of not only decreased iron concentration but altered iron trafficking within erythroblasts. The cytosolic chaperone Poly(rC)-binding protein 1 (PCBP1) delivers iron to ferritin [54,55] with evidence from *Pcbp1* knockout mice, with microcytosis and anemia, that iron delivery to ferritin is required for normal erythropoiesis [55]. In addition, PCBP2 is also required for ferritin complex formation [54]. Conversely, an autophagic process to extract iron from the ferritin core is mediated by nuclear receptor coactivator 4 (NCOA4), a selective cargo receptor for autophagic ferritin turn-over, critical for regulation of intracellular iron availability [56,57]. In iron replete states, PCBP1 and PCBP2 expression is enhanced while NCOA4 is targeted to the proteasome for degradation [58,59]. Our results demonstrate that mRNA expression of both *Pcbp1* (Figure 8A-8C) and *Pcbp2* (Supplementary Figure 15A-15C) in sorted bone marrow ProE, BasoE, and PolyE is elevated in MDS mice and does not return to control WT levels in DFP-treated MDS mice. Conversely, mRNA expression of both *Pcbp1* (Figure 8D) and *Pcbp2* (Supplementary Figure 15D) in sorted bone marrow OrthoE is decreased in MDS relative to WT mice, normalized in DFP-treated MDS mice.

**Figure 8.**
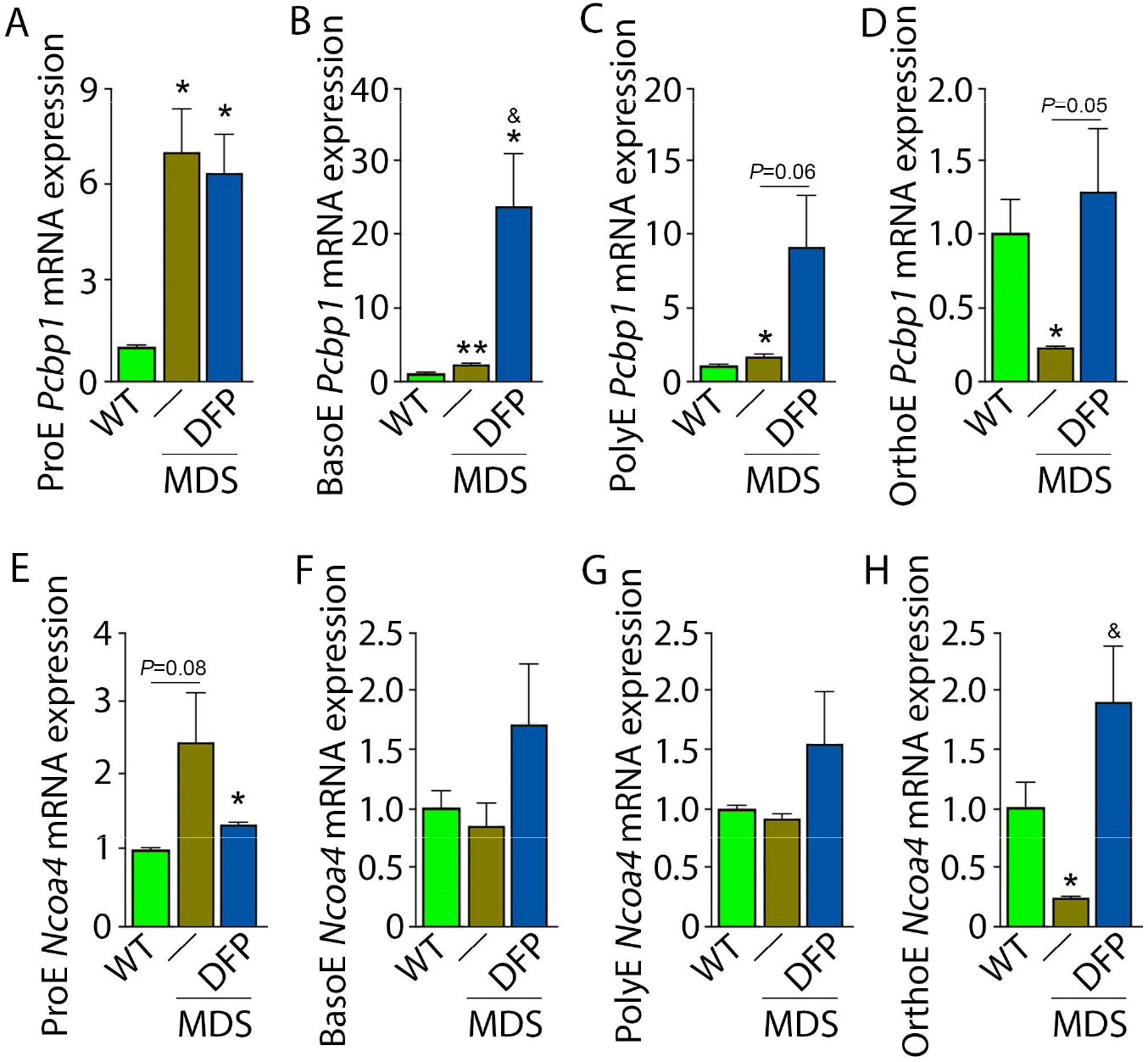
DFP alters expression of iron chaperones in MDS erythroblasts. Sorted bone marrow *Pcbp1* mRNA expression is increased in ProE **(a)**, BasoE **(b)**, and PolyE **(c)** from MDS relative to WT mice and remains high or is further elevated in DFP-treated relative to untreated MDS mice. **(d)** Sorted bone marrow *Pcbp1* mRN A expression is decreased in OrthoE from MDS relative to WT mice, normalized in DFP-treated MDS mice. Sorted bone marrow mRNA expression of *Ncoa4* in ProE **(e)** is borderline increased in MDS and remains elevated in DFP treated MDS mice. Sorted bone marrow mRNA expression of *Ncoa4* is unchanged in BasoE **(f)** and PolyE **(g)** from MDS or DFP-treated MDS relative to WT mice. Sorted bone marrow mRNA expression of *Ncoa4* in OrthoE **(h)** is decreased in MDS and normalized in DFP-treated MDS mice (n = 15-18 mice/group). \**P*<0.05vs. WT; ^&^*P*<0.05vs. MDS; Abbreviations: WT= wild type; MDS= myelodysplastic syndrome; DFP= deferiprone; *Pcbp1*= Poly(rC)-binding protein 1; *Ncoa4* = nuclear receptor coactivator 4; ProE = pro-erythroblasts; BasoE = basophilic erythroblasts; PolyE = polychromatophilic erythroblasts; OrthoE = orthochromatophi lie erythroblasts.

Furthermore, ProE mRNA expression of *Ncoa4* shows an increased trend in MDS mice and does not return to control WT levels in DFP-treated MDS mice (Figure 8E). While no differences in *Ncoa4* expression in bone marrow sorted BasoE and PolyE from WT, MDS, and DFP-treated MDS mice (Figure 8F and 8G), *Ncoa4* expression is suppressed in sorted bone marrow OrthoE in MDS mice, normalized in DFP-treated MDS mice (Figure 8H). Because of the role of NCOA4 in ferritinophagy and ferroptosis, we evaluate *Gpx4* expression in sorted bone marrow erythroblasts, demonstrating no differences between WT, MDS, and DFP-treated MDS mice (Supplementary Figure 16A-16D). These findings are consistent with expectations that high levels of iron flux through ferritin, high rates of ferritin turnover, and high rates of iron transfer to the mitochondria require elevated NCOA4 and PCBP1/2 levels [58] and provide preliminary evidence that movement of iron between sub-cellular compartments is altered in MDS erythroblasts, especially in early stages of terminal erythropoiesis, partially normalized by DFP.

### Increased Expression of Iron Metabolism Related Genes in MDS Patient Bone Marrow Stem and Progenitor Cells

Expression of iron metabolism related genes is compared in bone marrow derived CD34+ stem and progenitor cells from MDS patients (N=183) and healthy controls (N=17) as previously described [60]. As expected, *Tfr1, Epor, Gata1, Bcl2l1* (gene name for Bcl-Xl), and *Fam132b* expression is significantly increased in MDS patients relative to controls (Figure 9A-9E). In addition, while *Pcbp1* is unchanged and *Pcbp2* is borderline increased, *Ncoa4* is significantly increased in bone marrow stem and progenitor cells from MDS patients (Figure 9F-9H), enabling increased ferritin degradation in MDS erythroblasts. Finally, *Tfr2* expression is also significantly increased in MDS patients relative to controls (Figure 9I), confirming our results in MDS mice. Whether changes in iron sensing and trafficking within erythroblasts contribute to MDS pathophysiology and ineffective erythropoiesis more broadly is unexplored. Our findings demonstrate that NHD13 mice recapitulate pathophysiological changes in iron sensing and trafficking in erythroid progenitors from MDS patients.

**Figure 9.**
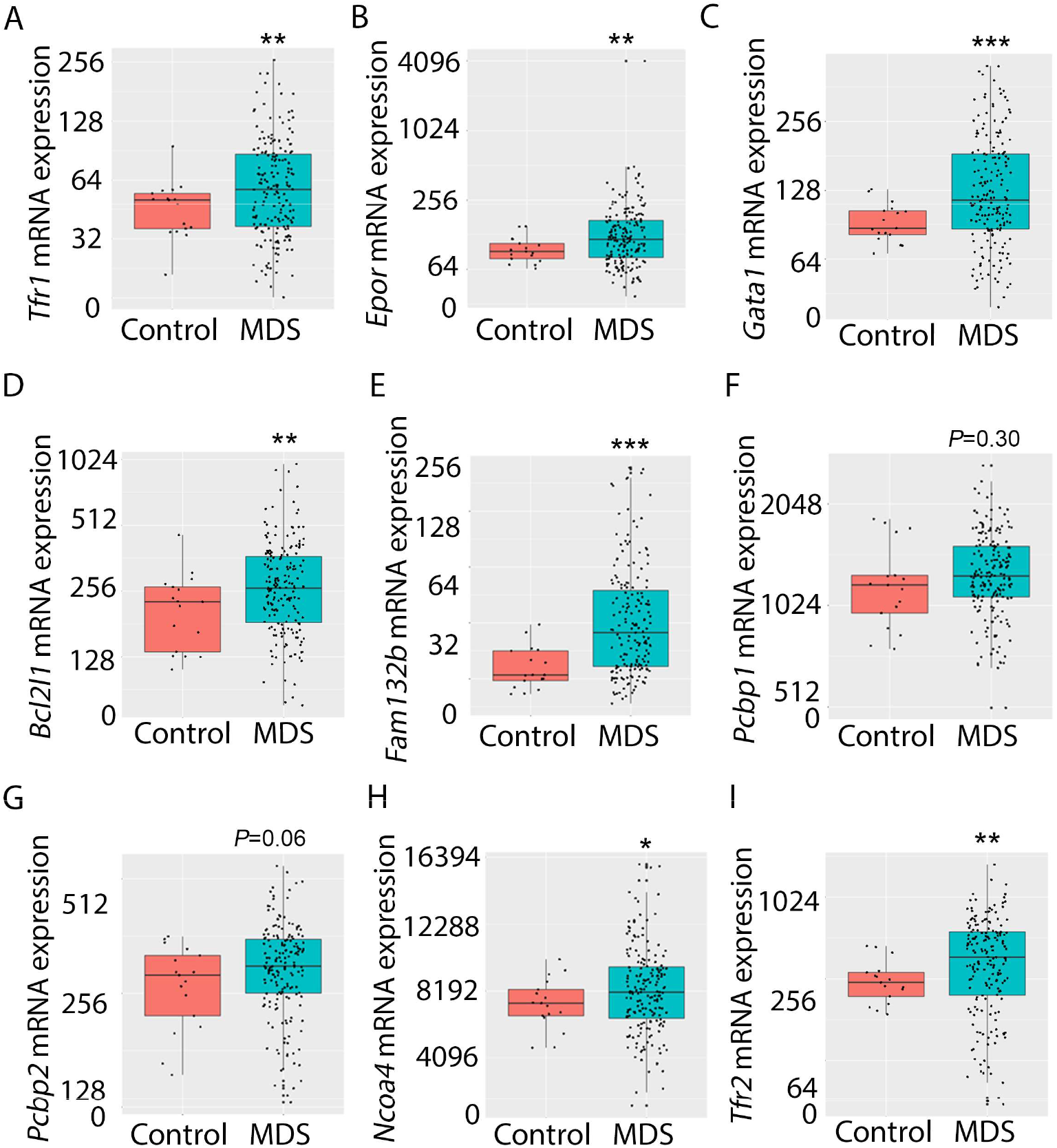
Increased expression of iron metabolism related genes in MDS patient bone marrow stem and progenitor cells. Increased expres sion of *Ttr1* **(a)**, *Epcr* **(b)**, *Gata1* **(c)**, *Bcl2l1* **(d)**, and *Fam132b* **(e)** is expected and validates the database. No difference in *Pcbp1* **(f)**, borderline increase in *Pcbp2* **(g)**, and statistically significantly increase *Ncoa4* **(h)** and *Tfr2* **(i)** are evident in MDS patients relative to controls, providing an important confirmation of the relevance of similar findings in MDS mice. * *P*<0.05, \*\**P*<0.01, *** *P*< 0.0001 vs. control; MDS = myelodysplastic syndrome; *Tfr1* and *Tfr2* = transferrin receptor 1 and 2; *Epor* = erythropoie tin receptor; *Be1211* = B-cell lymphoma 2-like protein 1 (gene name for Bol-Xl); *Fam132b* = erythroferrone; *Pcbp1* and *Pcbp2* = Poly(rC)-binding protein 1 and 2; *Ncoa4* = nuclear receptor coactivator 4.

## DISCUSSION

We show that MDS mice exhibit aberrant erythroblast iron trafficking and that iron chelation with DFP abrogates these changes to restore EPO-responsiveness (Figure 10). The effectiveness of DFP has previously been demonstrated in MDS patients [61]. These data raise the possibility that specifically targeting the iron sensing or iron trafficking machinery in erythroblasts may enable amelioration of ineffective erythropoiesis in addition to iron overload in MDS patients.

**Figure 10.**
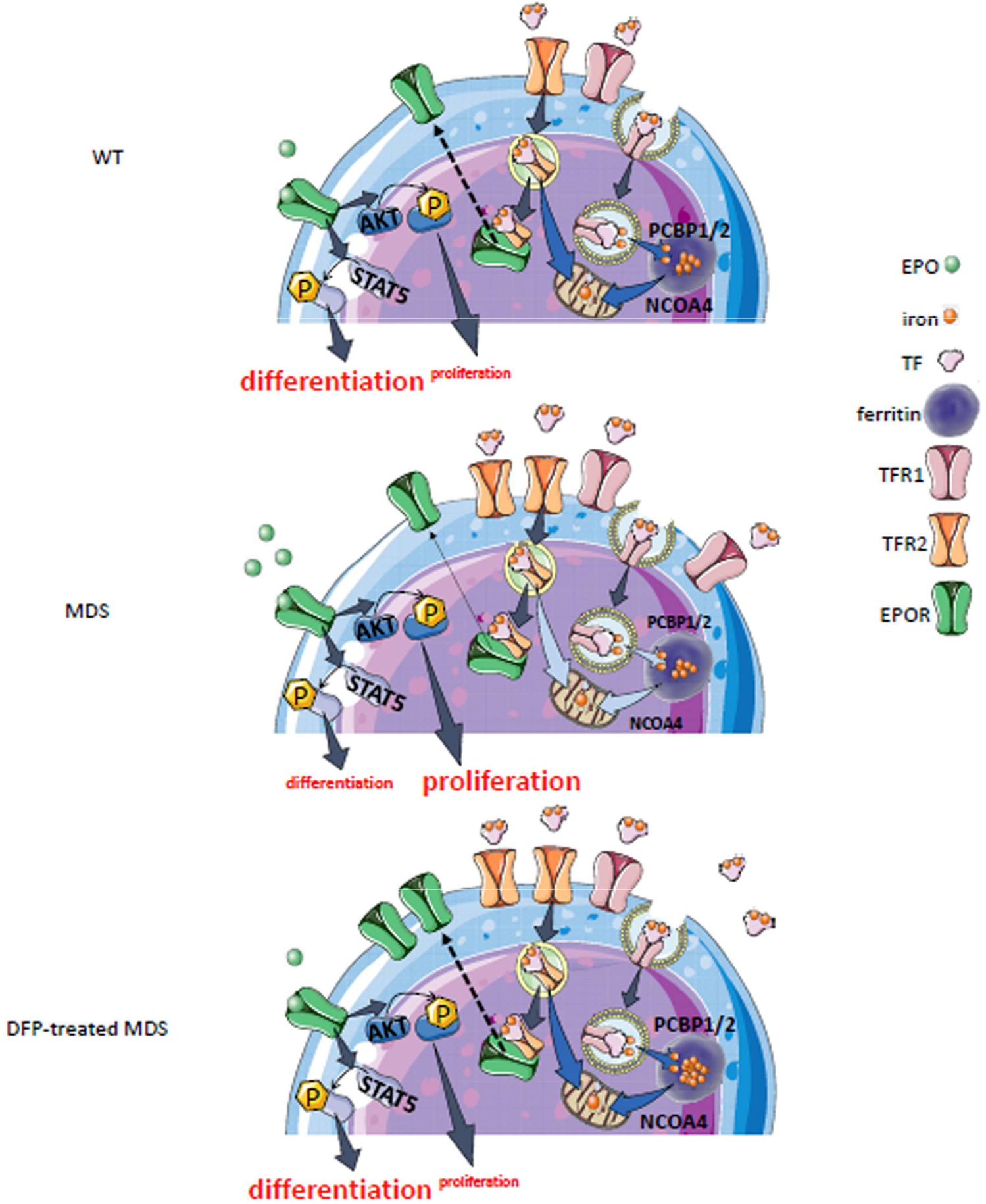
Putative function of erythrobIast iron trafficking in heaIth and in MDS before and after DFP treatment. Erythroblast iron uptake is mediated by TFR1 via endocytosis of clathrin coated pits while TFR2 is involved in iron sensing to coordinate iron supply with EPO responsiveness, modulating changes in signaling downstream of EPOR to alter erythroblast proliferation, differentiation, and apoptosis in response to physiological needs or pathophysiologic al conditions. In addition, erythroblast ferritin iron delivery is an obligatory step in erythropoiesis and chaperones, i.e. PCBP1 and PCBP2, deliver iron to ferritin while NCOA4 enable iron extraction from ferritin **(a)**. In conditions of expanded erythropoiesis, such as MDS **(b)**, more TFR1 results in increased iron uptake, resulting in increased TFR2; decreased EPOR, PCBP1 and NCOA4; yet no change in ferritin storage. We propose a modeI in which this pattern leads to altered iron delivery to the mitochondria, correlating with increased erythroblast survival, proliferation, and AKT signaling, but decreased erythroid differentiation. Treatment with DFP restores erythroblast iron trafficking with decreased TFR1, increased decreased PCBP1 and NCOA4, and increased ferritin concentration **(c)**. Abbreviations: WT = wild type; MDS = myelodysplastic syndrome; DFP = deferiprone; EPO = erythropoietin; EPOR = erythropoietin receptor; pSTAT5 = phosphotylated signal transducer and activator of transcription 5; pAKT = phosphorylated protein kinase B; TF = transferrin; TFR1 and TFR2 = transferrin receptor 1 and 2; PCBP 1/2 = Poly(rC)-binding protein 1 and 2; NCOA4 = nuclear receptor coactivator 4.

RBCs, the highest concentration among all cell types in circulation, are a product of erythroid precursor differentiation and enucleation, requiring 80% of the circulating iron for Hb synthesis [62]. Furthermore, how iron regulates erythropoiesis is incompletely understood and next to nothing is known about whether and how dysregulated iron metabolism contributes to the pathophysiology of ineffective erythropoiesis in MDS. The well-known, timely, abundant, and coordinated delivery of sufficient iron to erythroid precursors is accomplished via TFR1 to enable Hb production. TFR1 is both regulated by iron and enhanced by EPO-mediated signaling, evidence of the iron dependency in erythropoiesis. Furthermore, prior studies demonstrate that iron delivery to ferritin is absolutely required for normal erythropoiesis [55] and cytosolic chaperones PCBP1 and PCBP2 were recently identified as central to ferritin iron delivery in erythroblasts [54,55] (Figure 10). In addition, the process of ferritinophagy has recently been described, an autophagic process to extract iron from the ferritin core. NCOA4 is a selective cargo receptor for autophagic ferritin turn-over, critical for regulating intracellular iron availability for cellular function [56-59] (Figure 10). Finally, TFR2 expression has recently been identified in erythroid precursors [43,51-53, 63,64]. TFR2 is functionally involved in erythroid differentiation [53] via an interaction with EPOR [51] to modulate EPO responsiveness and possibly in shuttling iron to the mitochondria (Figure 10), but conflicting data findings underscore our incomplete understanding of down-stream effect of TFR2 in erythropoiesis. Taken together, movement of iron within erythroblasts is a complex multi-step process to handle a redox-active compound that is also required for the central task of erythropoiesis, i.e. Hb synthesis.

Our results demonstrate that iron trafficking is abnormal in MDS erythroblasts and restored in response to DFP. Specifically, increased TFR1 in MDS erythroblasts is expected to deliver more iron to erythroblast ferritin stores via increased PCBPs; more NCOA4 is expected to extract more iron from ferritin and more TFR2 to deliver more iron to mitochondria for Hb synthesis (Figure 10). Increased iron uptake by erythroblasts leads to a relatively normal amount of iron in the circulation despite systemic and parenchymal iron overload in MDS mice. Interestingly, erythroblast ferritin does not increase in MDS relative to WT mice; we speculate that this results from increased NCOA4-mediated iron release from ferritin and leads to an increase in iron’s redox activity and higher erythroblast ROS. Our findings suggest that, although regulation of late stage erythropoiesis remains incompletely understood, redox active iron accumulation may lead to a feedback downregulation in iron trafficking genes during erythroblast differentiation, consequently resulting in ineffective erythropoiesis.

Differences in early and later stage erythroblast can also be seen when comparing iron trafficking genes in MDS mice with that in stem and progenitor cells from MDS patients. Patient samples correlate with early stage erythroblasts in MDS mice in which *Tfr1, Epor, Tfr2*, and *Ncoa4* expression is increased, providing validation that altered iron trafficking is a relevant pathophysiological component in MDS patients. In further support, AML patients for example exhibit increased bone marrow TFR2 expression [65,66] and expression in blasts correlates with serum ferritin and overall prognosis [67]. Based on these finding, we propose that targeting iron trafficking in erythroblasts may be a viable therapeutic strategy in iron loading anemias, increasing erythroblast ferritin iron sequestration to protect later stage erythropoiesis from redox active effects of more labile forms of iron. For example, while NCOA4 is fundamental for iron supply during erythropoiesis, when iron is abundant, such as in MDS, other compensatory mechanisms are activated to prevent anemia [68], providing a rationale for targeting NCOA4 suppression to reverse ineffective erythropoiesis.

Lastly, although clearly increased ROS levels result in cellular damage, emerging evidence suggests that ROS is required for normal hematopoiesis, influencing stem cell migration, differentiation, cell cycle status, and self-renewal [69] such that hematopoietic stem cell self-renewal potential is associated with low ROS states while high ROS states are associated with differentiating hematopoietic stem cells. Further increased ROS leads to senescence and decreased ROS restores differentiation in those conditions [69]. In addition, ROS can activate JAK/STAT pathways [70] and therefore EPO-EPOR mediated cell growth and survival. Recent evidence indicates that EPO and iron are required for ROS generation in erythroblasts, and that ROS are necessary for terminal erythropoiesis [44] while unchecked ROS accumulation results in anemia [71-75]. Taken together, ROS generation is both critical and potentially toxic, requiring significant coordination during erythropoiesis with particular importance for mitigating increased ROS in conditions of ineffective erythropoiesis as in MDS. However, whether increased ROS resulting in apoptosis as the cause of ineffective erythropoiesis has never been definitely confirmed. We and others previously demonstrate that EPO downstream mechanisms are potent anti-apoptotic compensatory mechanisms and that reversal of apoptosis does not ameliorate ineffective erythropoiesis in β-thalassemia [76,77]. Our current results demonstrate borderline increased erythroblast apoptosis while ROS is significantly enhanced in MDS mice and no change in apoptosis while ROS is significantly decreased in DFP-treated MDS mice. These results uncouple apoptosis from ROS as the underlying cause of ineffective erythropoiesis.

In conclusion, despite FDA approval of iron chelators for use in iron overloaded MDS patients, they remain underutilized due to lack of biological insights into the deleterious effects of iron overload on disease pathophysiology in MDS. Our results demonstrate that DFP partially ameliorates ineffective erythropoiesis in MDS mice not only by reducing systemic iron overload, but also by altering iron trafficking within erythroblast and the sensitivity of erythroblasts to EPO, enhancing erythroblast differentiation. We further identify similar alterations in iron trafficking genes also in MDS patient bone marrow samples, validating our findings in MDS mice and anticipating a potential for translating these results. Taken together, these findings provide additional support for targeting erythroblast-specific iron sensing and trafficking to ameliorate ineffective erythropoiesis in MDS either with already available iron chelators, e.g. DFP, or novel therapies currently in development for diseases of ineffective erythropoiesis.

## MATERIALS AND METHODS

### Mice and treatment

C57BL6 (WT) and C57BL/6-Tg(Vav1-NUP98/HOXD13)G2Apla/J (NHD13) mice [29] were originally purchased from Jackson Laboratories (Bar Harbor, ME, USA). For simplicity, NHD13 mice are designated as “MDS mice” throughout the manuscript. All mice were bred and housed in the animal facility under Association for Assessment and Accreditation of Laboratory Animal Care guidelines. Experimental protocols were approved by the Institutional Animal Care and Use Committee. This well-established mouse model has been shown to recapitulate all key findings in human MDS, including blood cell dysplasia, peripheral blood cytopenias, ineffective hematopoiesis prior to transformation to acute leukemia, and a subset of mice progressing to acute leukemia at 14 months [29,30]. NHD13 mice on a C57BL/6 background were previously found to be clinical appropriate as an MDS model until at least 7 months [29,30]. As a consequence, we used age and gender matched 5-month old mice, at least 5 mice per group, treated with deferiprone (DFP; trade name Ferriprox™; chemical name 3-hydroxy-1,2-dimethylpyridin-4-one) at a concentration of 1.25 mg/mL in the drinking water for 4 weeks. DFP is an orally active iron chelator that binds iron in a 3:1 (DFP:iron) complex and undergoes renal clearance. Mice were euthanized for analysis at 6-months of age, as mice analyzed previously to study effects on ineffective erythropoiesis [28]. Therefore, all endpoints of interest were analyzed to compare DFP-treated MDS mice after 1 month of treatment with untreated MDS mice and WT controls.

### Peripheral blood analyses

Mice peripheral blood cell counting were analyzed by ProCyte Dx Hematology Analyzer. Serum mouse EPO (Quantikine, R&D Systems) was measured by enzyme-linked immunosorbent assay (ELISA) according to the manufacturer’s instructions. Integra 800 Automated Clinical Analyzer (Roche Diagnostics) was used to measure serum iron to transferrin iron-binding capacity (TIBC); serum transferrin saturation was measured as a ratio of serum iron to TIBC.

### Histology and immunohistochemistry

Immunohistochemical staining was performed using anti-TER119 antibodies (eBioscience, San Diego, CA), and counterstained with hematoxylin. Images were acquired on a Zeiss Axioskop2 microscope with an AxioCamHRC camera using Plan-Neofluar objectives ×20/0.5 and Axiovision software.

### Non-heme iron spectrophotometry

Quantification was performed via the Torrance and Bothwell method [78]. Briefly, specimens were digested overnight in acid solution at 65°C. A mixture of chromogen solution with acid extraction was incubated at room temperature for 10 minutes and the absorbance was measured at 540 nm by spectrophotometer (CLARIOstar plate reader, BMG Labtech).

### Quantitative real-time PCR

We prepared RNA from sorted bone marrow and liver samples using the RNeasy Kit (Qiagen) according to the manufacturer’s instructions. We synthesized cDNA using the High-Capacity cDNA Reverse Transcription Kit (Thermofisher). Primers are listed in Supplementary Table I. The qPCR was conducted by iQ™ SYBR®Green Supermix using the BioRad CFX96 Real Time PCR Thermal Cycler. Target gene mRNA concentration was normalized to *Gapdh* or *actin*.

### Western immunoblotting

Beads sorted CD45 negative bone marrow or liver cells were lysed in ice cold SDS page lysis buffer (2% SDS, 50 mM Tris-HCl, pH 7.4, 10 mM EDTA) with protease and phosphatase inhibitors. Twenty mg of heat–denatured protein was loaded onto a 10% gel, run, and transferred onto a 0.4 mm nitrocellulose membrane (Thermo Scientific). After blocking with 5% BSA in Tris– buffered saline with 1% Tween-20, the membranes were incubated with primary antibodies to signaling proteins (Supplement Table II) overnight at 4°C, washed, and incubated with the corresponding HRP–conjugated secondary antibodies at room temperature. Proteins were visualized using the ImageQuant LAS 4010 and quantified using Image J.

### Flow cytometric analysis and sorting

Bone marrow cells were processed as described previously [79] with minor modifications. Briefly, the cells were mechanically dissociated, blocked with rat anti–mouse CD16/CD32 (Fcγ III/II Receptor), incubated with anti-CD45 magnetic beads (Mylteni), and underwent magnetic separation using LS columns according to the manufacturer’s instructions (Miltenyi Biotec). Erythroid lineage-enriched CD45 negative cells were collected for further staining. Non-erythroid and necrotic cells were excluded using anti-CD45 (BD Pharmigen), anti-CD11b, and anti-Gr1 (APC-Cy7) (Tonbo, Biosciences) antibodies. Cells were incubated with anti-mouse TER119 PE-Cy7 (BioLegend) and CD44-APC (Tonbo, Biosciences) to identify and delineate progressive stages of erythroblast differentiation (Supplementary Figure 17). Once erythroblasts were delineated by TER119, CD44 and forward scatter, CD71-PE (Biolegend) was used to evaluate changes in erythroblast membrane TFR1. ROS quantification in erythroblasts was performed using immunostaining for ROS (Invitrogen) as per manufacturer’s instructions. To evaluate apoptosis, cells were stained for activated caspase 3/7 kit (Invitrogen); 7-amino-actinomycin D (7AAD, BD Pharmingen) was added to exclude dead cells. Cells were analyzed within 1 hour of staining using BD FACSDiva Version 6.1.2 software on a FACSCanto flow cytometer (Becton Dickinson). The gating strategy was as previous described [79]. Erythroid differentiation was quantified by analyzing the fraction of each stage of terminal erythropoiesis relative to all erythroblasts in each bone marrow sample. In addition, to individually evaluate gene expression in erythroblasts at different maturation stages, bone marrow cell underwent sorting on a BD FACSAria™ III (BD Biosciences). Finally, enucleation was assessed as described previously [80].

### Gene expression in MDS patient database

Gene expression data from 183 MDS CD34+ samples and 17 controls were obtained from GEO (GSE19429) as previously described [60].

### Statistical analyses

All data are reported as mean ± standard error of the mean (SEM). We performed analysis for statistically significant differences with the 2-tailed student paired *t* test.

## ACKNOWLEDGEMENTS

We sincerely appreciate the perpetual energy and devotion of the late Professor Eliezer Rachmilewitz to the importance of understanding and treating iron overload in MDS and dedicate this manuscript to his memory. We are grateful for funding from CSL Behring (King of Prussia, PA) (to Y.Z.G.) and the Italian Government Fellowship (to M.F.) for support of early work that served as the foundation of this study and ApoPharma (Chiesi) for supporting the experiments presented here.

---

**Supplementary Table I.**
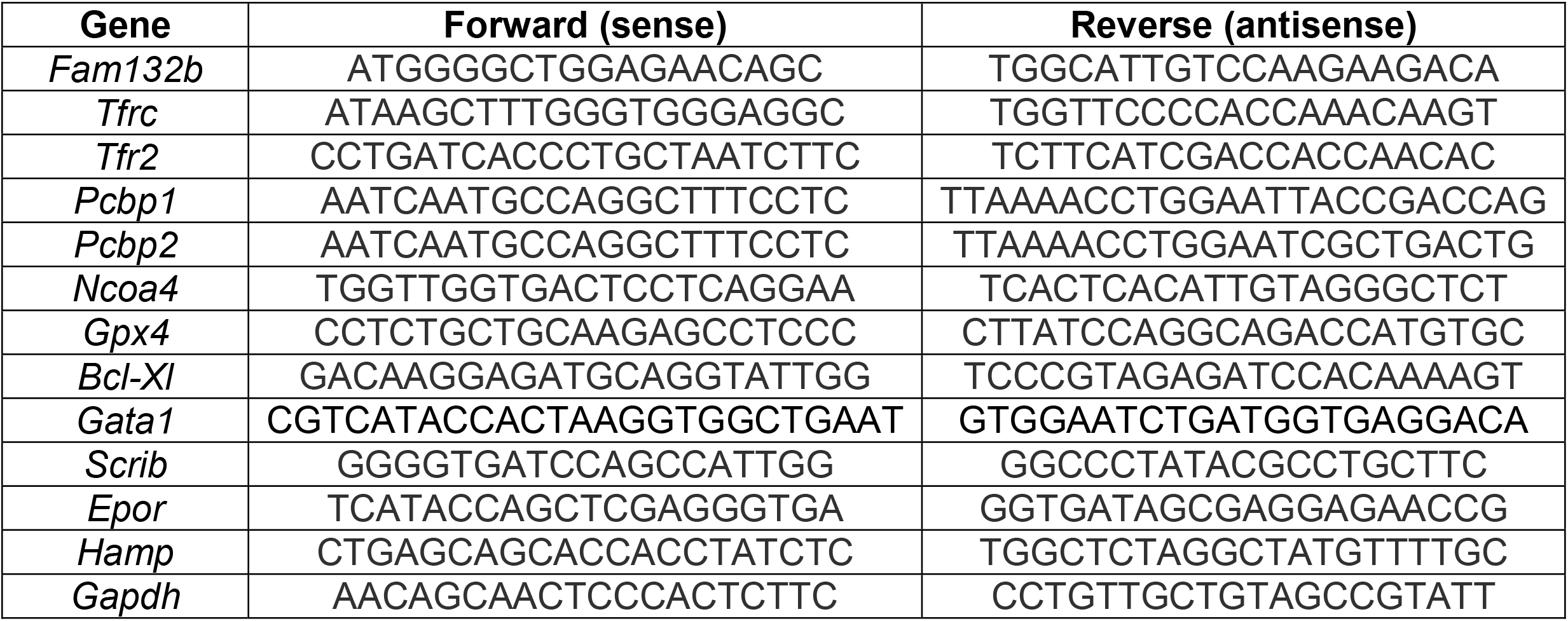

**Supplementary Table II.**

| **Antibody** | **Company** | **Catalog #** |
| --- | --- | --- |
| Beta-actin | ThermoFisher | MA515452 |
| GAPDH | Cell Signaling Technology | 97166 |
| Phospho-STAT3 | Cell Signaling Technology | 9131 |
| STAT3 | Cell Signaling Technology | 12640 |
| Phospho-STAT5 | Cell Signaling Technology | 9359 |
| STAT5 | Cell Signaling Technology | 94205 |
| Phospho-AKT | Cell Signaling Technology | 4058 |
| AKT | Cell Signaling Technology | 4691 |
| FTH1 | Cell Signaling Technology | 4393 |
| TFR2 | Abcam | ab80194 |

**Supplementary Figure 1.**
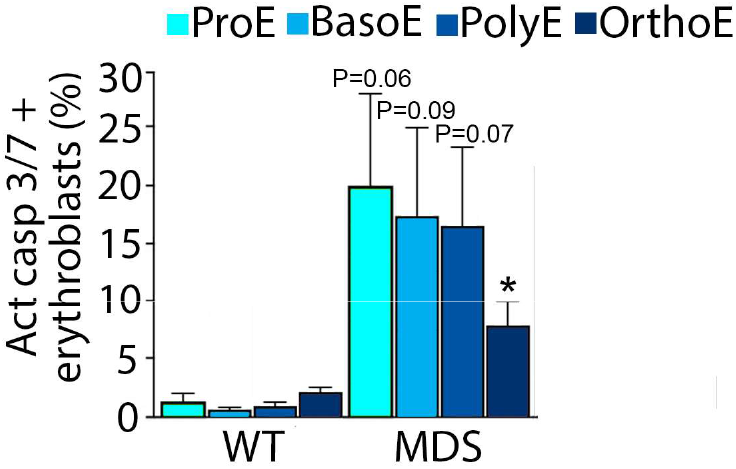
Effect of DFP on erythroblast apoptosis in MDS mice. Bone marrow erythmblasts were isolated using flow gating strategy and apoptosis was measured using activated caspase 3/7. Erythroblast apoptosis is borderline elevated in ProE, BasoE, and PolyE and significantly elevated in OrthoE fmm MDS mice (n = 7-11 mice/group). * *P*<0.05 vs. WT; WT = wild type; MDS = myelodysplastic syndrome; ProE = pro-erythroblasts; BasoE = basophilic erythroblasts; PolyE = polychromatophilic erythroblasts; OrthoE = orthochromatophilic erythroblasts; Act casp 3/7 = activated caspase 3 and 7.

**supplementary Figure 2.**
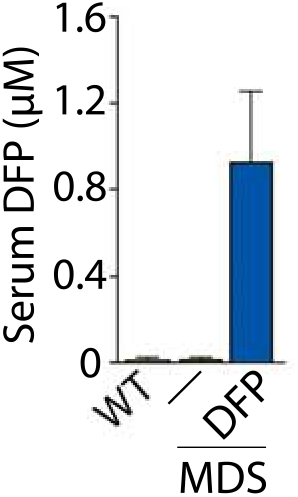
Quantification of serum DFP concentration in DFP-treated WT and MDS mice. Serum DFP concentration is measurable in DFP-treated MDS mice. (n = 4-6 mice/group). WT= wild type; MDS= myelod ysplastic syndrome; DFP = deferiprone.

**Supplementary Figure 3.**
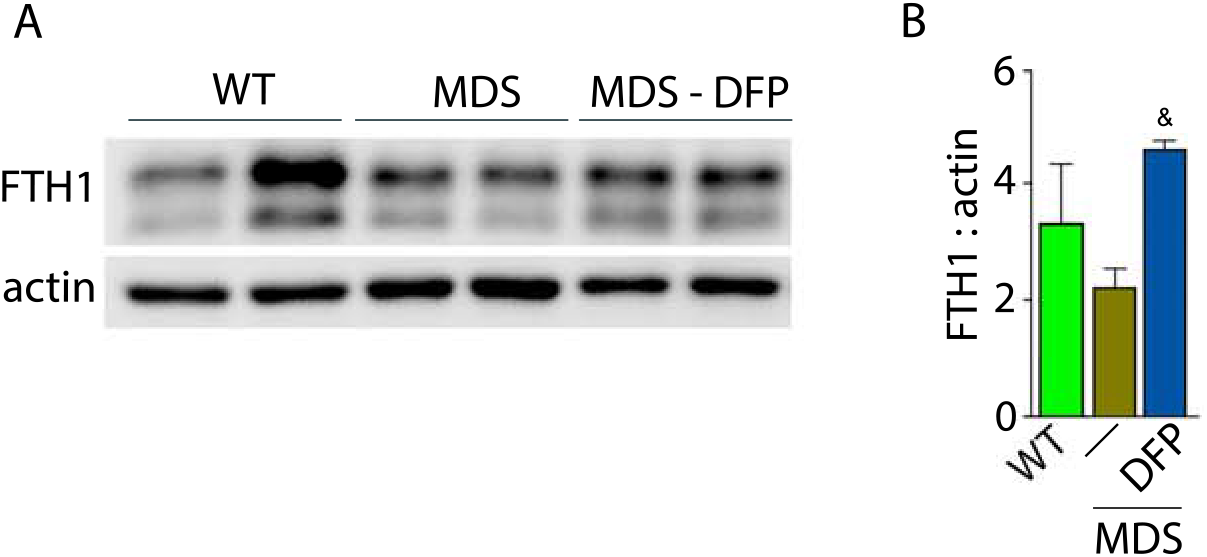
Bone marrow erythroblast ferritin is increased in DFP-treated MOS mice. **(a)** Western blot of bone marrow CD45 negative cell protein extracts demonstrate no difference in FTH1 between WT and MDS mice, increased to normal levels in DFP-treated MDS mice; gel is quantified in **(b)** (n = 2 mice / group, experiments repeated twice). WT = wild type; MDS = myelodysplastic syndrome; DFP = deferiprone; FTH1 = ferritin heavy chain.

**Supplementary Figure 4.**
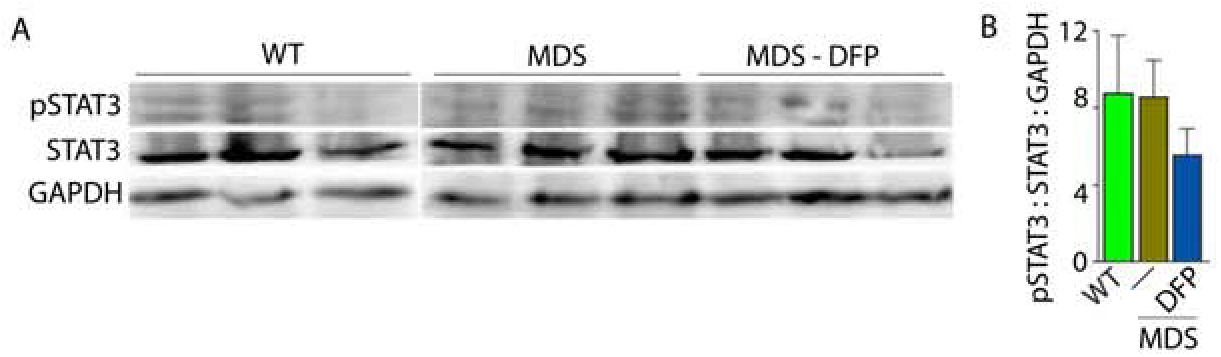
Effects of DFP on the liver STAT3 expression in M DS mice. **(a)** Western blot of liver protein extracts demonstrate no differences in STAT3 signaling, quantified in **(b)**, demonstrating no change In the inflammatory signaling pathway to hepcidln expression between WT, MDS, and DFP-treated MDS mice (n = 3 mice/ group). WT= wild type; MDS = myelodysplastlc syndrome: DFP = deferiprone; pSTAT3 = phospho1ylated signal transducer and activator of transcription 3; GAPDH = glyceraldehyde 3-phasphate dehydrogenase.

**Supplementary Figure 5.**
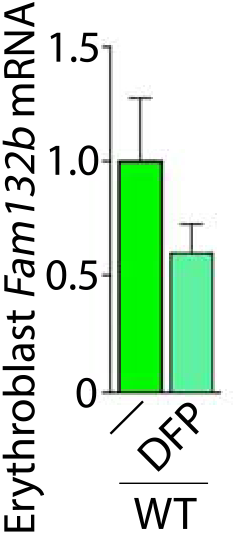
Effect of DFP on erythroblast *Fam132b* expression in WT mice. Sorted bone marrow erythroblast *Fam132b* mRNA expression between WT and DFP-treated WT mice (n = 5-15 mice/group). \**P*<0.05 vs. WT DFP; WT= wild type; DFP = deferiprone; Fam132b = erythrorerrone.

**Supplementary Figure 6.**
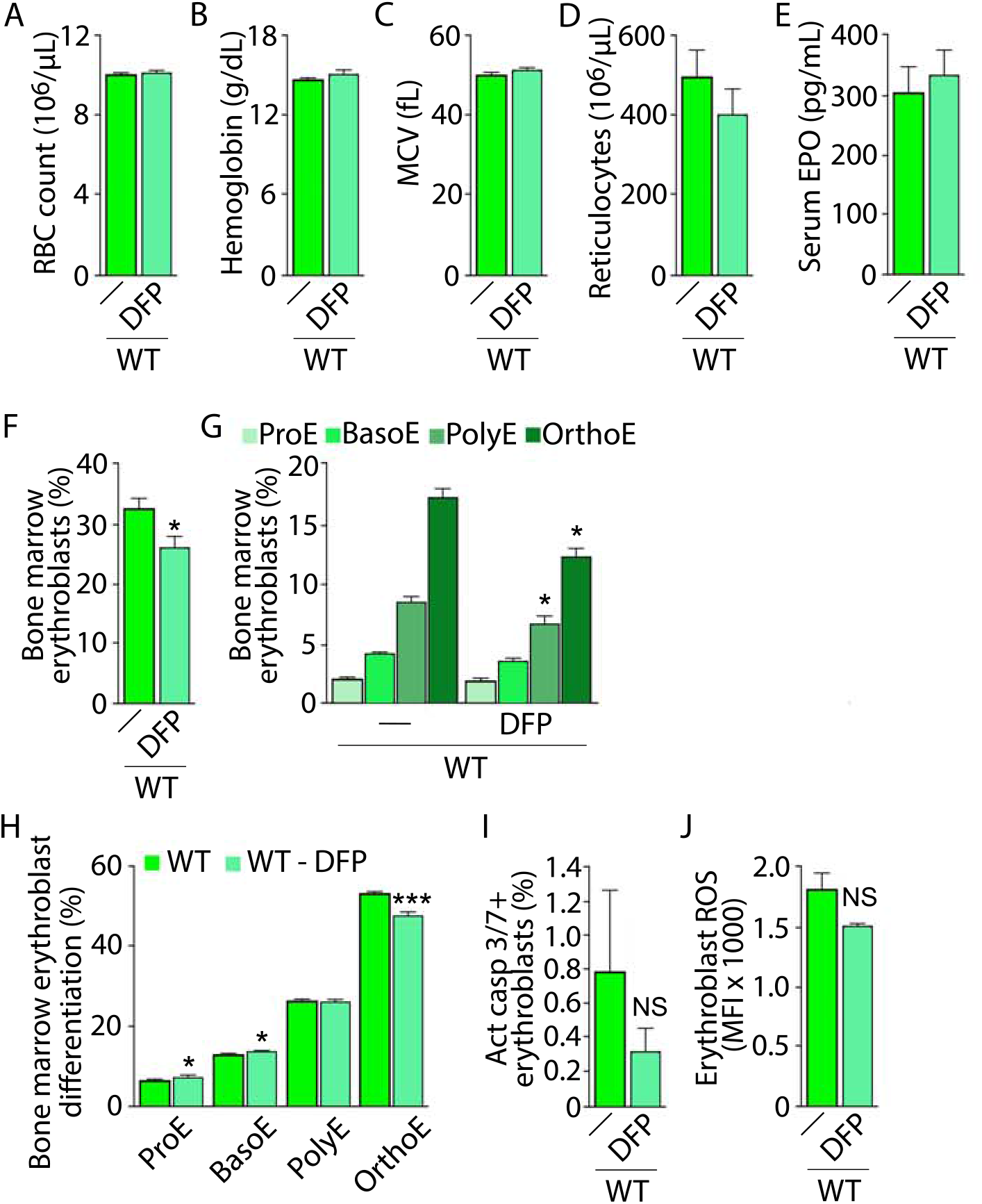
Effect of DFP on erythropoiesis In WT mice. No differences are observed In RBC count **(a)**, hemoglobln **(b)**, MCV **(c)**, reticuocytes **(d)**, or serum EPO concentration **(e)**. Bone marrow erythroblast fraction significantly decreased **(f)** particularly as a consequence of fewer PolyE and OrthoE **(g)** and consistent with decreased differentiation **(h)** in DFP-treatedWT mice. Erythroblast apoptosls, as measured by activaed caspase 3 and 7 (I), and ROS (i) are unchanged in DFP-treated relative to untreated WT mice (n = 5-11 mice/group). * *P*<0.05 vs. WT DFP; WT= wild type: DFP = deferiprone: RBC = red blood cell; MCV = mean corpuscular volume: EPO = erythropoietln; ProE = pro-erythroblasts; BasoE = basophilic erythroblasts; PolyE = polychro matophlllc erythroblasts; OrthoE = orthochromatophlllc erythroblasts; Act casp 3/7 = activated caspase 3 and 7; ROS = reactive oxygen species; NS = net significant.

**Supplementary Figure 7.**
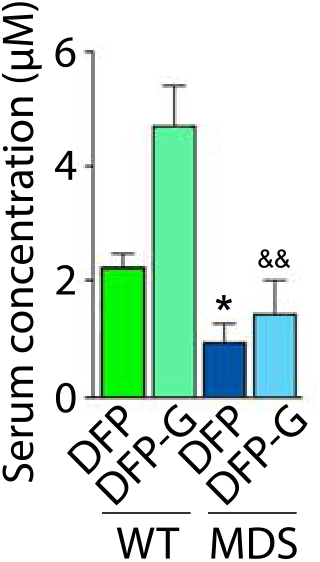
Quantification of serum DFP-glucuronlde concentration In DFP treated WT and MDS mice. serum coocentration rt DFP and or its metabolite, DFP-G, is significartly lower in DFP-treated MOS relGCive to DFP-treated WT mice (n = 4-6 mice/gro) * *P*<0.05 vs. WT DFP: ^&&^*P*<0.01 vs. WT DFP-G; WT = wild type: MDS = myelodysplastlc syndrome; DFP = deferiprone; DFP-G = DFP-glucuronide.

**Supplementary Figure 8.**
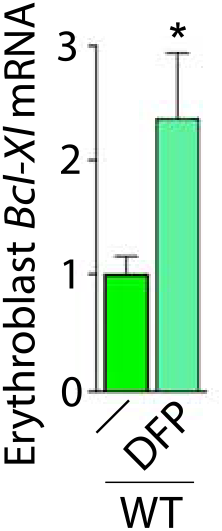
Effect of DFP on erythroblast *Bel-XI* expression in WT mice. Sorted bone marrow erythroblast *Bel-XI* mRNA expression between WT and OFP-treated WT mice (n=5-15 mice/group). * *P*<0.05 vs. WT DFP; WT =wild type; DFP =deferiprone; Bcl-XI=8-cell lymphoma-extra large.

**Supplementary Figure 9.**
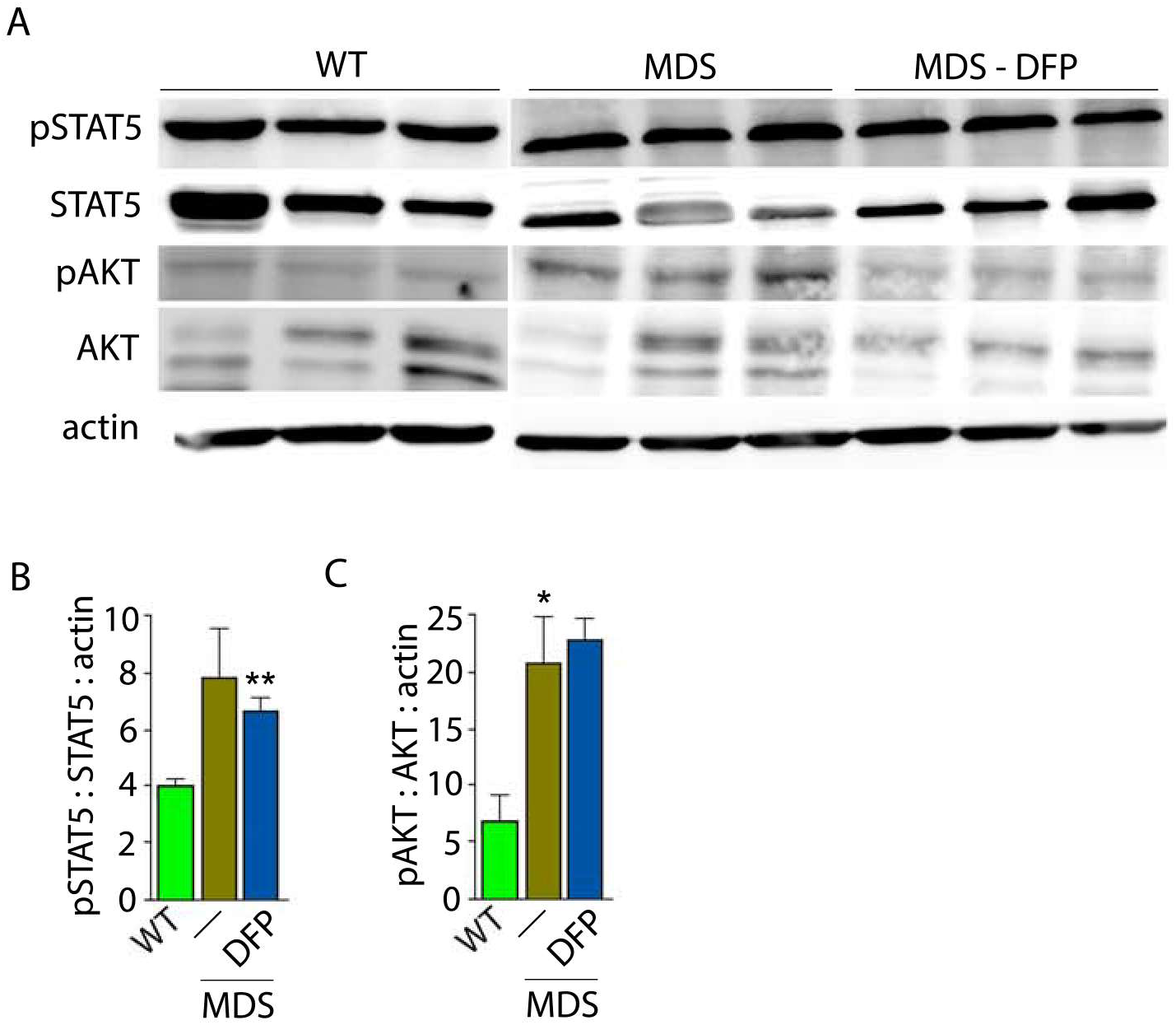
Nuances of signaling downstream of EPO in bone marrow erythroblasts from WT, MDS, and DFP-treated MDS mice. **(a)** Western blot of bone marrow CD45 negative cell protein extracts demonstrate no differences in STAT5 signaling between WT and MDS mice and no normalization in DFP-treated MDS mice, quantified in **(b)**. Similarly, AKT signaling is elevated in MDS relative to WT and not normalized in DFP-treated MDS mice, quantified in **(c)**, demonstrating no change in these signaling pathways in response to DFP in MDS mice (n = 3 mice / group). WT = wild type; MDS = myelodysplastic syndrome; DFP = deferiprone; pSTAT5 = phosphorylated signal transducer and activator of transcription 5; pAKT = phosphorylated protein kinase B. * *P*<0.05vs. WT; \*\**P*<0.01 VS. WT.

**Supplementary Figure 10.**
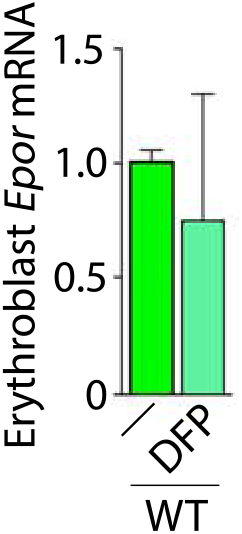
Effect of DFP on erythroblast *Epor* expression in WT mice. Sorted bone marrow erythroblast EpormRNA expression between WT and DFP-treated WT mice (n = 5-15 mice/group). * *P*<0.05 vs. WT DFP; WT = wild type; DFP = deferiprone; Epor = erythropoietin receptor.

**Supplementary Figure 11.**
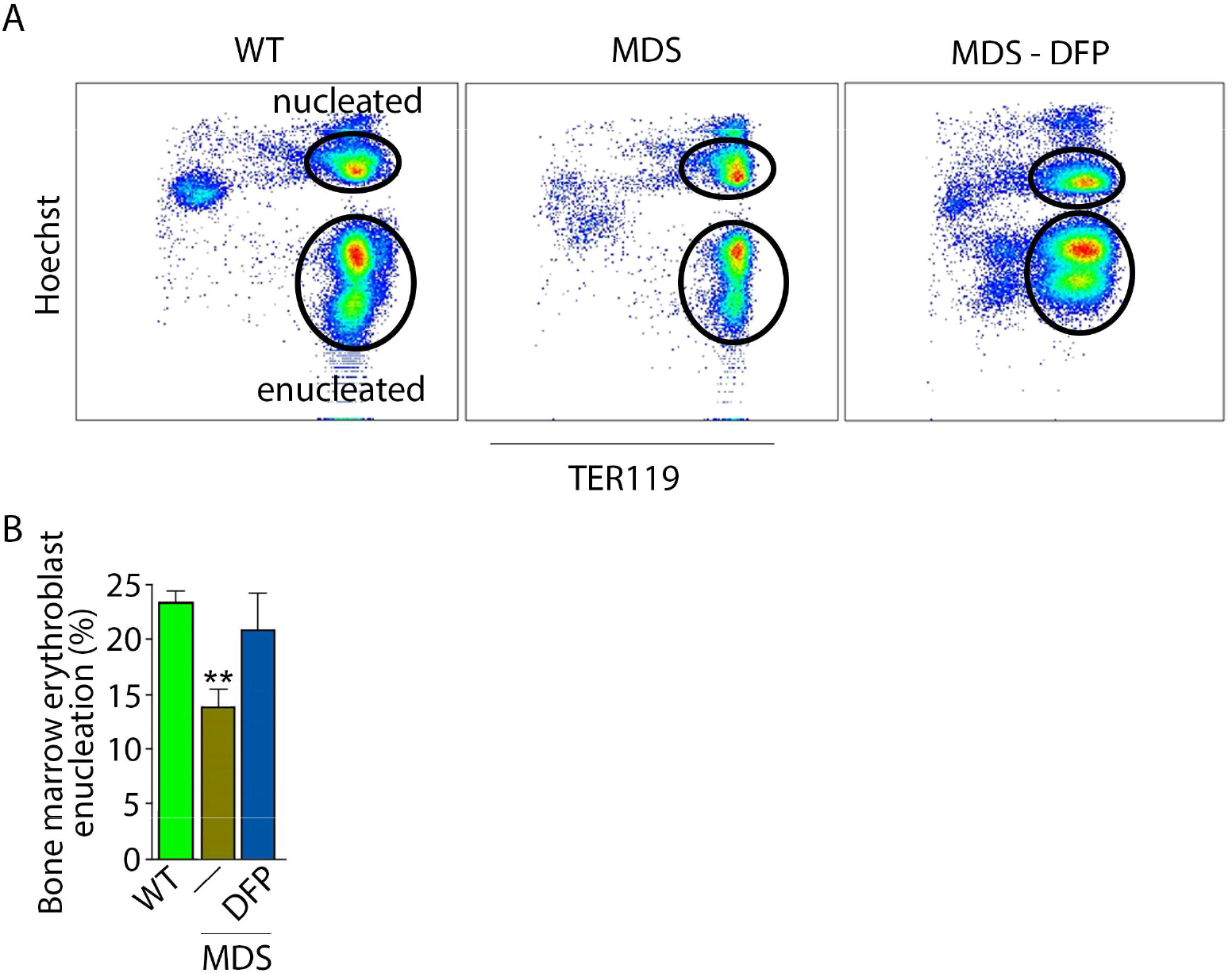
Detective enucleation in MDS erythroblasts is normalized by DFP. **(a)** OFP restores erythroblast enucleation (n= 6-7 mice/group) in MDS mouse b one marrow, quantified in **(b)** as the fraction of enucleated relative to total of nucleated and enucleated erythroblasts, using Hoechst staining in TER119 positives. \*\**P*<0.1 vs. WT; Abbreviations: WT = wild type; MDS = myeloctys plastic syndrome; DFP = deferiprone.

**Supplementary Figure 12.**
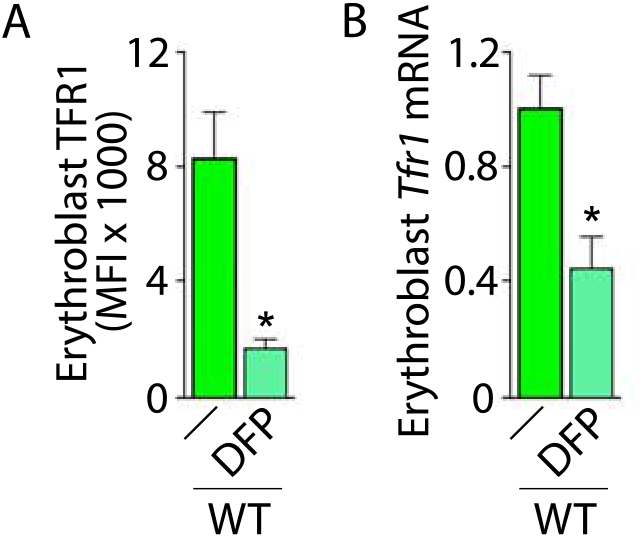
Effect of DFP on erythrobla.st *Tfr1* and TFR1 expression in WT mice. **(a)** Sorted bone marrow erythroblast *Tfr1* mRNA expression between WT and DFP-treated WT mice (n = 5-12 mice/group). **(b)** Membrane TFR1 on bone marrow eryttlroblasts from WT and DFP-treated WT mice (n = 5-11 mice/group). * *P*<0.05 vs. WT DFP; WT = wild type; DFP = deferiprone: Tfr1 = transferrin receptor 1: MFl = mean fluorescence index.

**Supplementary Figure 13.**
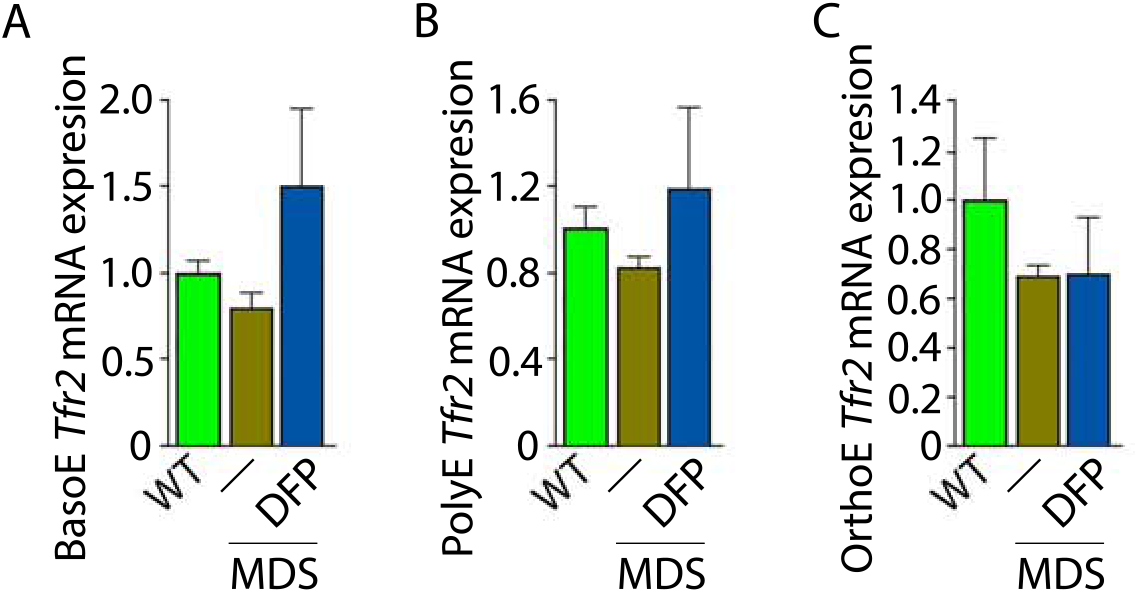
Erythroblast Tfr2 mRNA expression In WT, MDS, and DFP treated MDS mice. No differences are observed In *Tfr2* mRNA expression In sorted BasoE **(a)**, PolyE **(b)**, and OrthoE **(c)** In **WT**, MDS, and DFP-treated MDS mice (n: 5-15 mice ***/*** group), WT = wild type; MDS = myelodysplastic syndrome: DFP = defenprone; Tfr2 transferrin receptor 2; BasoE = basophilic erythroblasts; PolyE : polychromatophilic erythroblasts; OrthoE = orthochromatophilic erythroblasts.

**supplementary Figure 14.**
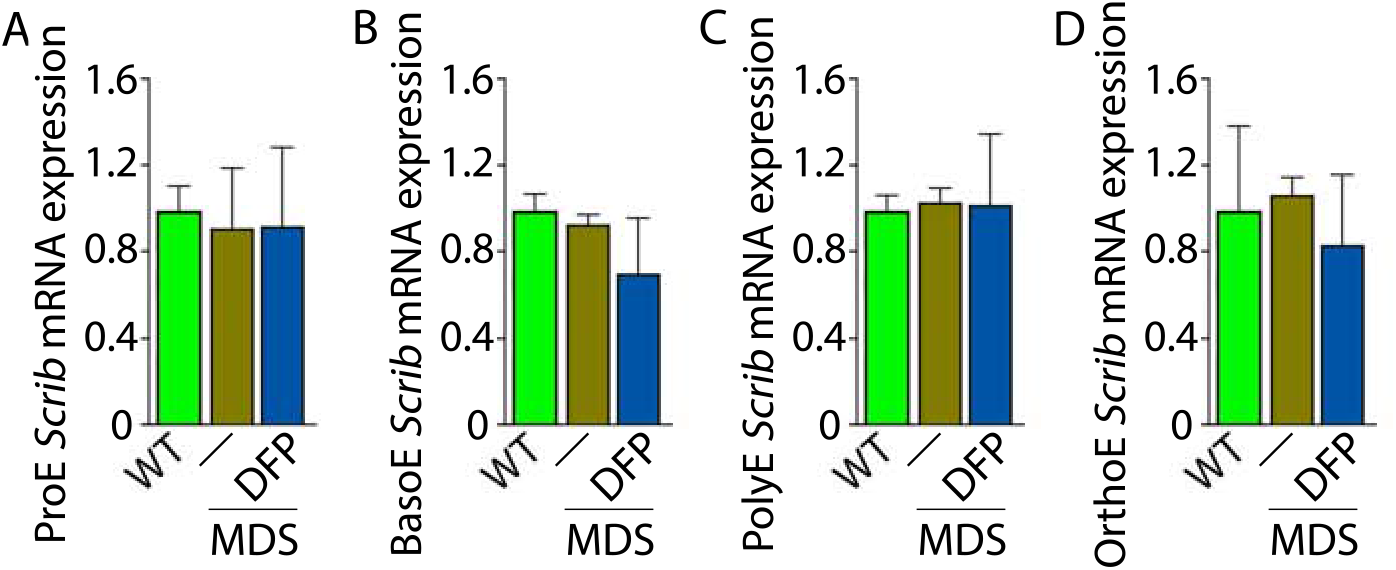
Erythroblast *scrib* mRNA expression in WT, MDS, and DFP· treated MDS mice. No differences In *Scrib* mRNA expression are observed In sorted bone marrow ProE **(a)**, BasoE **(b)**, PolyE **(c)**, and OrthoE **(d)** between WT, MDS, and DFP-treated MDS mice (n 15-18 mice / group). WT wild type; MDS myelodysptastlc syndrome; DFP deferiprone; ProE = pro-erythroblasts; BasoE = basophilic erythroblasts; PolyE = polychromatophilic erythroblasts; OrthoE = orthochromatophilic erythroblasts; Scrib = Scribble.

**Supplementary Figure 15.**
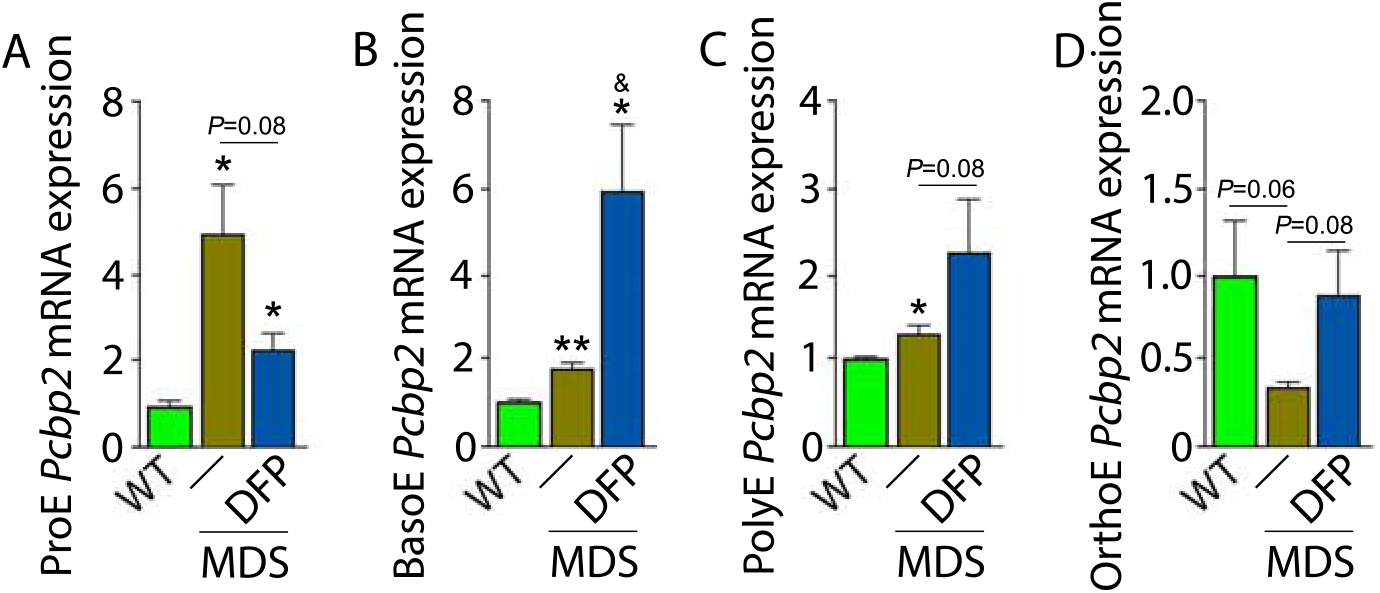
Erythroblast *Pcbp2* mRNA expression in WT, MDS, and DFP· treated MDS mice. *Pctp2* mRNA expression changes parallel those of *Pcbp1* in sorted bone marrow ProE **(a)**, BasoE **(b)**, PolyE **(c)**, and OrthoE **(d)** between WT, MDS, and DFP-treated MDS mice (n = 15-18 mice / group). WT= wild type; MDS = myelodysplastlc syndrome; DFP = deferiprone: ProE = pro-erythroblasts; BasoE = basophilic erythroblasts; PolyE = polychromatophilic erythroblasts; OrthoE = orthochromatophllic erythroblasts; Pcbp2 = Poly(rC) binding protein 2.

**Supplementary Figure 16.**
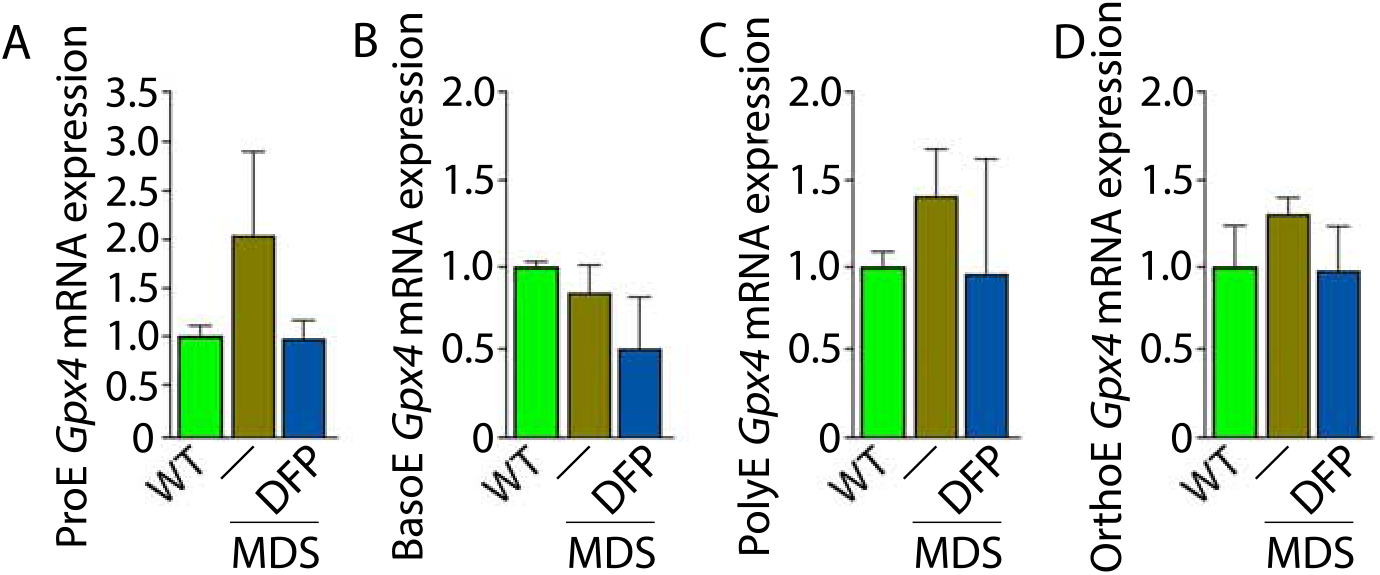
Erythroblast *Gpx4* mRNA expression in WT, MDS, and DFP-treated MDS mice. No differences in *Gpx4* mRNA expression are observed in sorted bone marrow ProE **(a)**, BasoE **(b)**, PolyE **(c)**, and OthoE **(d)** between WT, MDS, and DFP-treated MDS mice (n = 15-18 mice / group). WT= wild type; MDS = myelodysplastic syndrome; DFP = deferiprone; ProE = pro-erythroblasts; BasoE = basophilic erythroblasts; PolyE = polychrornatophilic erythroblasts; OrthoE = orthochromatophilic erythroblasts; GPX4 = glutatnione peroxidase 4.

**supplementary Figure 17.**
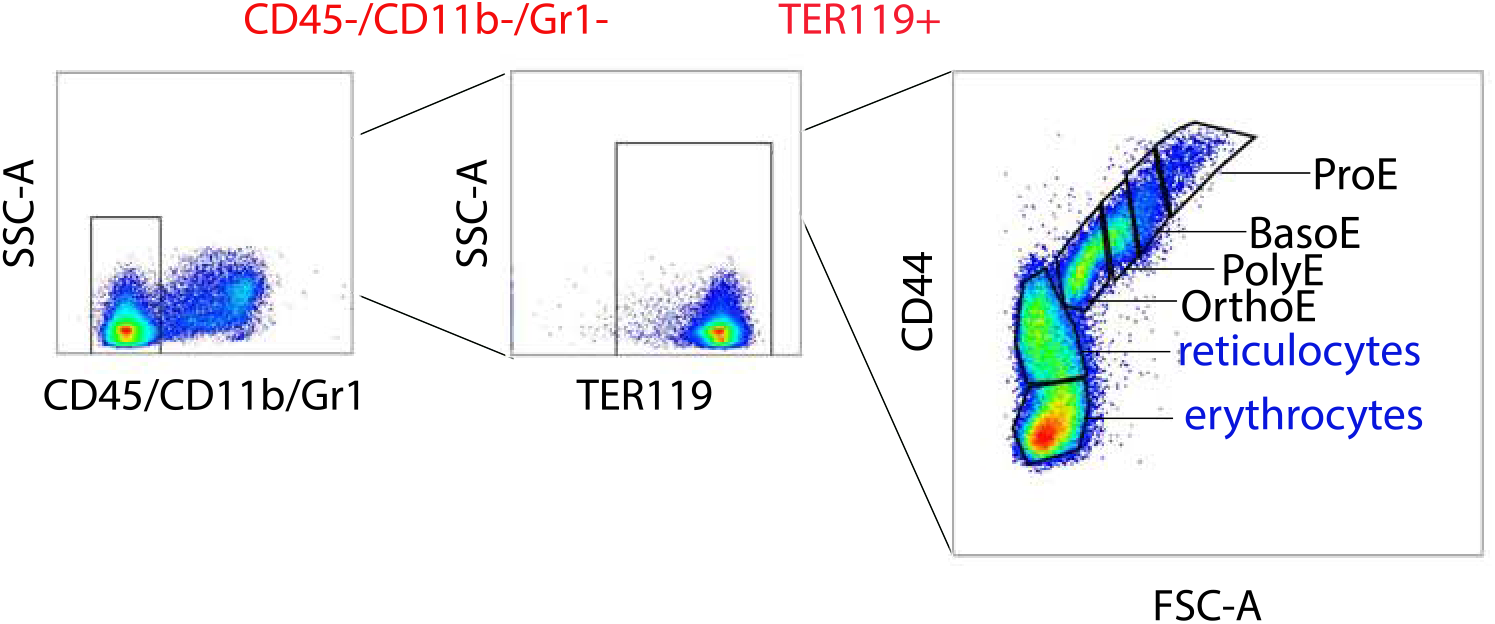
Gating strategy for delineating erythroblasts in mouse bone marrow. Mouse bone marrow is flushed from the femur, processed to remove debris, and filtered to collect CD45 negative flow through cells. After staining with all required antibodies, CD45-/CD11b-/Gr1-cells are gated (red) and further evaluated using TER119 and side scatter (SSC) to select all TER119+ erythroid lineage cells (red). These cells are then anatyzed using forward scatter (FSC) and CD44 to gate pro-erythroblasts (ProE), basophilic erythroblasts (BasoE), polychromatophillc erythroblasts (PolyE), and orthochromatophilic erythroblasts (OrthoE) as progressive stages of terminal erythropoiesis and exclude reticulocytes and enucleated erythrocytes (blue).

